# PLMFit : Benchmarking Transfer Learning with Protein Language Models for Protein Engineering

**DOI:** 10.1101/2025.01.15.633186

**Authors:** Thomas Bikias, Evangelos Stamkopoulos, Sai. T. Reddy

## Abstract

Protein language models (PLMs) have emerged as a useful resource for protein engineering applications. Transfer learning (TL) leverages pre-trained parameters to extract features to train machine learning models or adjust the weights of PLMs for novel tasks via fine-tuning through back-propagation. TL methods have shown potential for enhancing protein predictions performance when paired with PLMs, however there is a notable lack of comparative analyses that benchmark TL methods applied to state-of-the-art PLMs, identify optimal strategies for transferring knowledge and determine the most suitable approach for specific tasks. Here, we report PLMFit, a benchmarking study that combines, three state-of-the-art PLMs (ESM2, ProGen2, ProteinBert), with three TL methods (feature extraction, low-rank adaptation, bottleneck adapters) for five protein engineering datasets. We conducted over >3,150 *in silico* experiments, altering PLM sizes and layers, TL hyperparameters and different training procedures. Our experiments reveal three key findings: (i) utilizing a partial fraction of PLM for TL does not detrimentally impact performance, (ii) the choice between feature extraction and fine-tuning is primarily dictated by the amount and diversity of data and (iii) fine-tuning is most effective when generalization is necessary and only limited data is available. We provide PLMFit as an open-source software package, serving as a valuable resource for the scientific community to facilitate the feature extraction and fine-tuning of PLMs for various applications.

**ONE SENTENCE SUMMARY:** PLMFit is a comparative analysis aimed at identifying the most effective strategies for transfer knowledge from protein language models by benchmarking fine-tuning techniques on a range of protein engineering tasks.

## INTRODUCTION

Protein language models (PLMs) are becoming a valuable tool in computational biology with applications such as protein structure and function prediction, design and engineering^1–3^. Leveraging transformer-based architectures^4^ and inspired by large language models (e.g. ChatGPT^5^, Llama^6^, Falcon^7^), PLMs are trained on a large corpora of unlabeled protein sequences and are able to generate sequences of amino acids representing proteins with a high likelihood of folding, expression and biological function. During this process, known as pre-training, multi-layered PLMs capture evolutionary^8^ and structural^9^ dependencies between amino acids by attempting to either reconstruct a corrupted sequence (i.e. masked language modeling) or predict the next residue (i.e. token) given the previous as context (i.e. autoregressive language modeling). Acquired knowledge is stored in the weights of the different PLM layers and can transform the input sequence into an information rich representation: protein sequence embeddings. As an alternative to encoding only amino acid sequence information (i.e., one-hot encoding, OHE) or including evolutionary information extracted from multiple sequence alignments (i.e., BLOSUM matrix)^10^, PLM embeddings can be used as input features to train, typically shallow, machine learning (ML) models [i.e., artificial neural networks (ANNs) or convolutional neural networks (CNNs)] to solve a wide variety of protein characterization and engineering tasks^11–15^.

Transfer learning (TL) leverages pre-trained parameters to train or adjust a different model to a novel task; TL can broadly be divided, based on the adaptation of the pre-weights or not, into two categories: feature extraction (FE) and fine-tuning (FT). In the context of PLMs, FE employs the retrieval of pre-trained weights from a PLM layer, which converts protein residues into evolutionary informed features^16^ that can be used as inputs for training novel models. On the contrary, FT includes the joint optimization of PLM (or PLM fraction) weights with an untrained network (i.e. downstream head) using labeled data. To increase generative and downstream performance, PLMs continue to scale to a larger number of parameters and training sets, which poses technical challenges for implementing FT. Arbitrary retraining of PLMs, especially recently established large models (e.g., ESM-3^17^), is computationally infeasible and can cause catastrophic forgetting of previously acquired knowledge^18^. Strategies adapted from natural language processing (NLP) may be able to mitigate these issues. For example, parameter-efficient fine-tuning (PEFT) is able to adapt pre-trained models to a new domain with minimal adjustments to the original parameters. These techniques focus on optimizing only a small subset of newly added parameters, rather than retraining the entire network, allowing lower resource consumption and maintaining or improving performance on the specific task. Commonly used methods include low-rank adaptation (LoRA)^19^, which involves adding low-rank trainable matrices in parallel with the transformer layers and adapter modules^20^, which uses the injection of small neural network modules in between each layer of the pre-trained model.

Several studies propose pairing of TL techniques with PLMs to extract meaningful representation of proteins^3,21,22^, this is accomplished mainly by using the embeddings extracted from the last layer of the PLM as input features to train novel ML models and by investigating the effect of PEFT for applications in protein engineering ^23–25^. However, it is still unclear in which setups the exploitation of PLMs has a guaranteed benefit, as baseline models trained with simple encoding schemes (i.e., OHE) have been shown to overperform PLM-based methods in relevant protein engineering tasks^26^. Moreover, choosing the most appropriate TL methods is not straightforward. Multiple factors require calibration in order to optimally retrieve the stored information (e.g. extraction layer, downstream head architecture, FT hyperparameters and more). Additionally, TL is impacted by the amount, diversity, and quality of training data as well as access to hardware resources (i.e., memory and number of GPUs). Recent studies have evaluated the effectiveness of PEFT methods applied to PLMs for addressing biology-related tasks^27^, or attempt to identify which specific layers might be most beneficial for embeddings extraction^22,28^. However, analysis of PLM layer-specific performance including both FE and PEFT methodologies has yet to be reported. Additionally, comparative studies with baseline models and very large PLMs (> 5 billion parameters) are missing.

Here, we report PLMFit, a comparative analysis that benchmarks TL methods applied on state-of-the-art PLMs and explores the best strategies for transferring stored knowledge from these models, identifying the most suitable approaches for specific protein engineering challenges. We apply PLMFit on six publicly available datasets corresponding to different protein properties and fitness prediction and classification tasks. Using labeled training datasets of protein sequences and functions, we evaluate >3,000 TL configurations by varying the following: (i) PLM architectures and sizes, (ii) layers of PLMs, (iii) FT hyperparameters and (iv) different training procedures. Our experiments reveal three key findings: (i) utilizing a fraction of PLM for transfer learning does not detrimentally impact performance, (ii) the choice between feature extraction and fine-tuning is primarily dictated by the amount and diversity of data and (iii) fine-tuning is most effective when generalization is necessary and only limited data is available. PLMFit thus offers a practical guide on how to best leverage pre-trained models and TL based on the nature of the protein engineering task and available computational resources. We provide PLMfit as an open-access software platform to seamlessly apply TL on user data using PLMs. (Figure 1)

**Figure 1.**
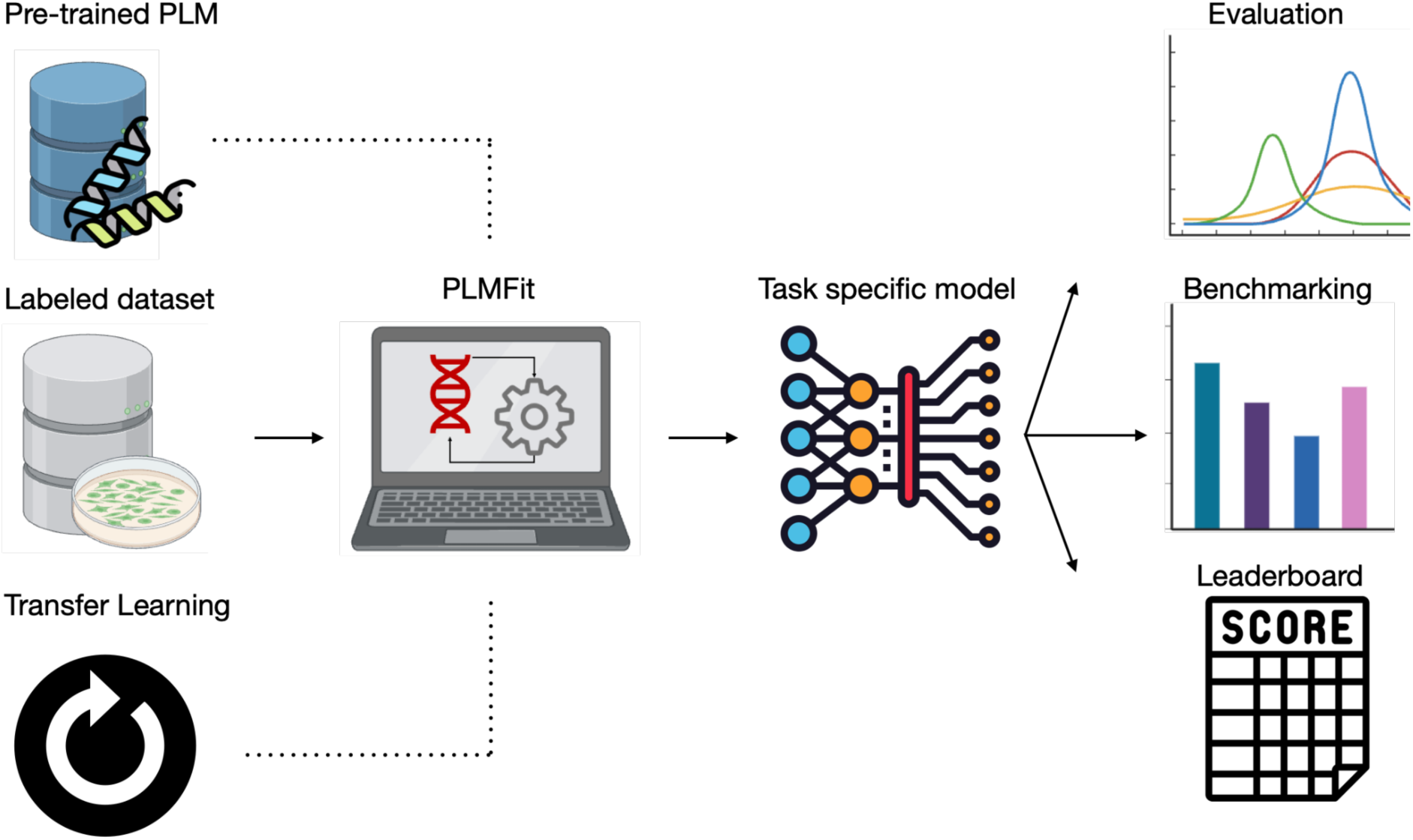
Overview of the PLMFit benchmarking analysis. PLMFit platform task-specific TL workflow. A pre-trained PLM can be paired with TL technique (FT or FE) with a labeled dataset. The resulting task-specific model is applied to a downstream task and benchmarked against alternative methods. PLM: Protein Language Models; TL: Transfer Learning; FE: Feature Extraction; LoRA: Low Rank Adaptation

## RESULTS

### Utilizing datasets that represent a broad range of protein engineering tasks

To establish benchmarks for the various TL techniques, we utilized publicly available repositories of protein sequence and function datasets that correspond to different types of tasks and levels of complexity, attempting to cover common use cases of protein engineering. In the context of this study, we consider different splits (i.e. how the training and test data are separated) for each task and we refer to a configuration as the combination of a PLM and a specific set of TL methods and hyperparameters applied on a specific training and evaluation procedure. In the protein engineering and design field, training datasets consist of two main types: (i) protein sequence mutational landscape, which refers to the range of possible mutations that can occur, including substitutions, insertions, or deletions and the edit distance (ED) is the number of mutations a given variant has relative to a reference sequence [i.e., wild-type (WT)], where the number of mutations and distribution of residues remains relatively consistent and (ii) sequences of proteins from different families with a more diverse distribution of residues. For the first category, we utilized four fitness regression datasets parsed from the fitness landscape inference for proteins (FLIP) repository^15^ and two protein-protein interaction (binding) classification datasets ^29,30^ (Table 1). In this context, fitness refers to a protein’s ability to perform a function across different taxonomic groups and is influenced by multiple environmental factors, including stability, binding affinity, enrichment, foldability, and catalytic activity. For example, in antibody discovery, antibodies are experimentally mutated in various regions to investigate the change in binding affinity to a specific antigen^31^. For the second category (i.e. diverse datasets), we used the *Meltome* dataset from the FLIP repository, which includes melting temperature measurements for a diverse range of proteins and the secondary structure prediction (i.e., SS3) dataset^24^, which involves predicting three-state secondary structures (helix, strand, coil) for sequences drawn from multiple protein families. Thus, protein sequences *in SS3* present considerable variation in residue composition and arrangement. We suggest that the complexity of each task is determined by both the quantity and diversity of the data available for training (i.e., training splits), as well as the nature of the testing data on which the prediction will be applied. Throughout the manuscript, we refer to a task using the dataset name and the split separated by a hyphen “-” (e.g., *AAV-sampled*). Thus, two different tasks can refer to the same dataset but differentiate in complexity because of the training split (i.e, *AAV-sampled* and *AAV-one vs. rest).* We define tasks as simple, such as *AAV-sampled* and *GB1-three vs. rest* due to the similarity in the distribution of the training and evaluation sets, which both consist of protein sequences with similar EDs from WT. In contrast, we define tasks as complex, such as *one vs. rest* splits, as they involve limited training data that often consist of single point mutation variants (ED = 1) but require predictions on variants with a higher number of mutations (i.e. ED ≥ 2). We consider the *Meltome-mixed* and the *SS3-sampled* task to be complex due to the high variability in the sequences used. Additional details about the datasets and tasks used can be found in “Datasets & Methods” section “Datasets and downstream tasks”.

**Table 1.**
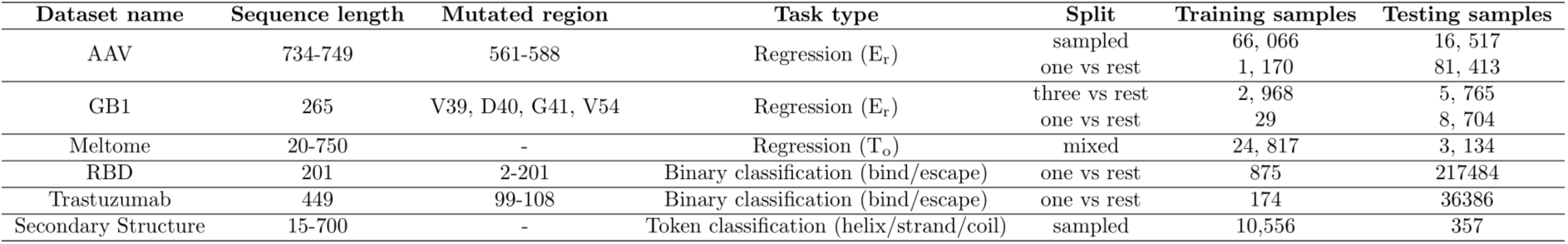
Summary of datasets and splits used for each dataset utilized in this study.

| Dataset name | Sequence length | Mutated region | Task type | Split | Training samples | Testing samples |
| --- | --- | --- | --- | --- | --- | --- |
| AAV | 734-749 | 561-588 | Regression ($E_r$) | sampled<br>one vs rest | 66,066<br>1,170 | 16,517<br>81,413 |
| GB1 | 265 | V39, D40, G41, V54 | Regression ($E_r$) | three vs rest<br>one vs rest | 2,968<br>29 | 5,765<br>8,704 |
| Meltome | 20-750 | - | Regression ($T_o$) | mixed | 24,817 | 3,134 |
| RBD | 201 | 2-201 | Binary classification (bind/escape) | one vs rest | 875 | 217484 |
| Trastuzumab | 449 | 99-108 | Binary classification (bind/escape) | one vs rest | 174 | 36386 |
| Secondary Structure | 15-700 | - | Token classification (helix/strand/coil) | sampled | 10,556 | 357 |

### Transfer Learning using a fraction of Protein Language Models impacts tasks’ performance

We evaluated three TL approaches: FE, LoRA and adapters (Figure 2) for their performance across four different tasks (i.e., *AAV-sampled, GB1-three vs. rest, Meltome-mixed, SS3-sampled*) and across different layers (first layer only, 25%, 50%, 75%, 100%) from three PLM families (ESM, ProGen and ProteinBERT) (Figure 3) (see Datasets & Methods). Particularly across the tasks *AAV-sampled* and *GB1-three vs. rest*, we observed that performance plateaued when 25% of layers were used for all three TL methods, after which there were minimal gains or even drops in performance. This suggests that pre-trained parameters stored in the first quarter of a foundational model may be more suitable when used for tasks that include protein sequences with similar distribution (i.e. variants of a WT) (Figure 3Ai-ii, Figure 3Bi-ii, Figure 3Ci-ii). Using these early parameters provides a simpler representation that focuses on general features, making them better suited for capturing subtle differences among closely related sequences. Thus, the final layers of a PLM are not always the best choice for transfer learning. This pattern is consistent across all TL configurations that involve sequences within the same ED distribution between the training and the testing set. For tasks involving diverse protein sequences such as *Meltome-mixed* (Figure 3A-Ciii) and *SS3-sampled* (Figure 3A-Civ), incremental performance benefits are exhibited when deeper layers are targeted for TL, indicating that the task’s complexity and sequence variability benefits from richer and more comprehensive representations from the deeper layers of PLMs. This hints that, for highly diverse and complex tasks, leveraging the full depth of larger models can provide a distinct advantage, as they capture more nuanced features and dependencies across diverse protein families. Performance differences among the various PLMs, despite their differing architectures and sizes, are relatively marginal. While larger models like ProGen2-xlarge and ESM2-15B perform slightly better in most setups, shallower models such as ProteinBERT and ProGen2-small achieve comparable results even on complex tasks. This suggests that the model size and architecture do not always dictate performance when effective TL techniques are applied. The choice of TL method and the number of layers used appears to have a greater influence on performance, except in the case of the *SS3-sampled*, where larger ESM models (i.e., ESM2-3B and ESM2-15B) demonstrate a notable improvement in performance (Figure 3B-Civ).

**Figure 2.**
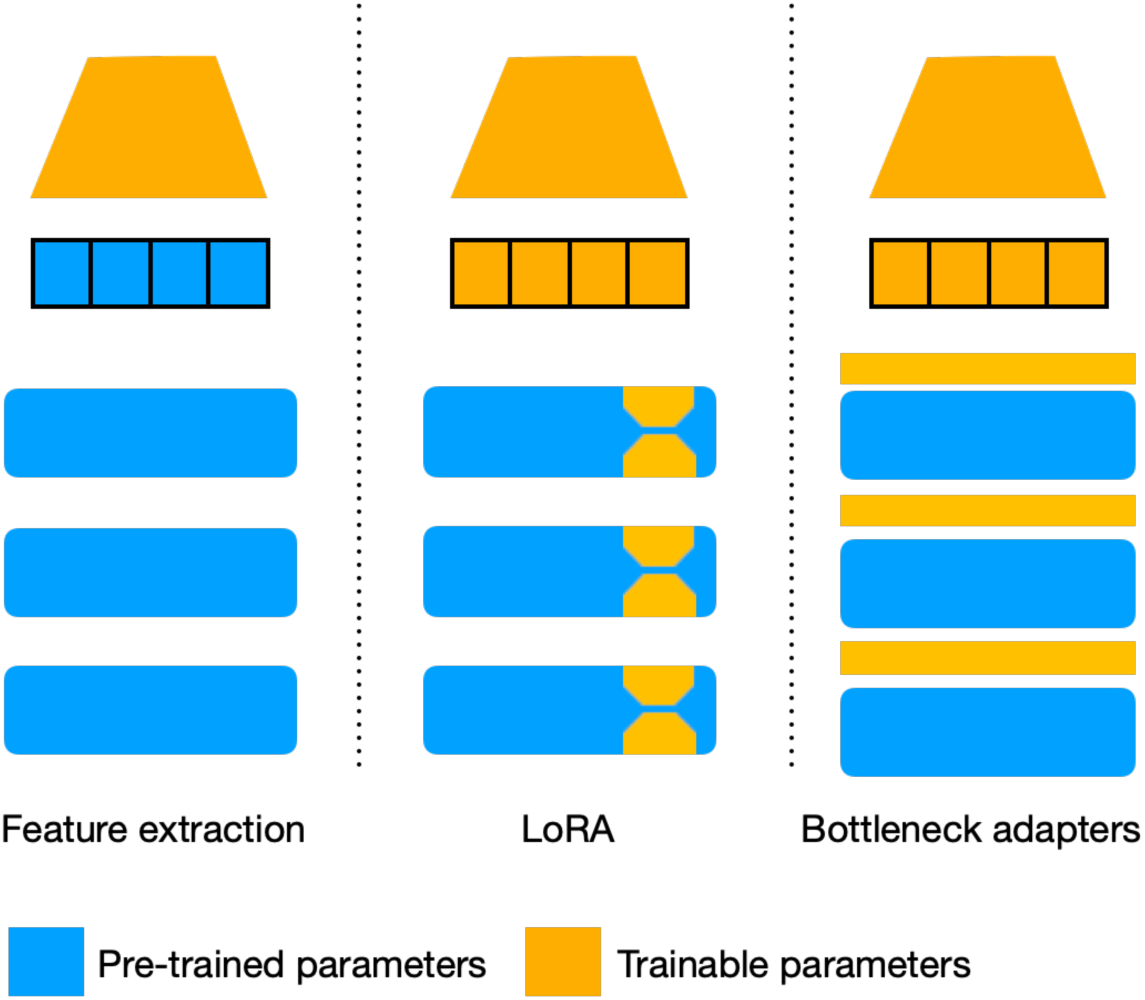
Transfer learning strategies used with protein language models. FE involves keeping the pre-trained model parameters (blue boxes) frozen while using them to extract features. LoRA is parameter efficient FT technique that introduces trainable parameters (orange) into low-rank matrices within the layers to adapt the pre-trained model for a specific task. Similarly, Bottleneck adapters inject trainable parameters within compact layers between the pre-trained parameters to adjust model outputs without full retraining. FE: Feature Extraction; LoRA: Low Rank Adaptation

**Figure 3.**
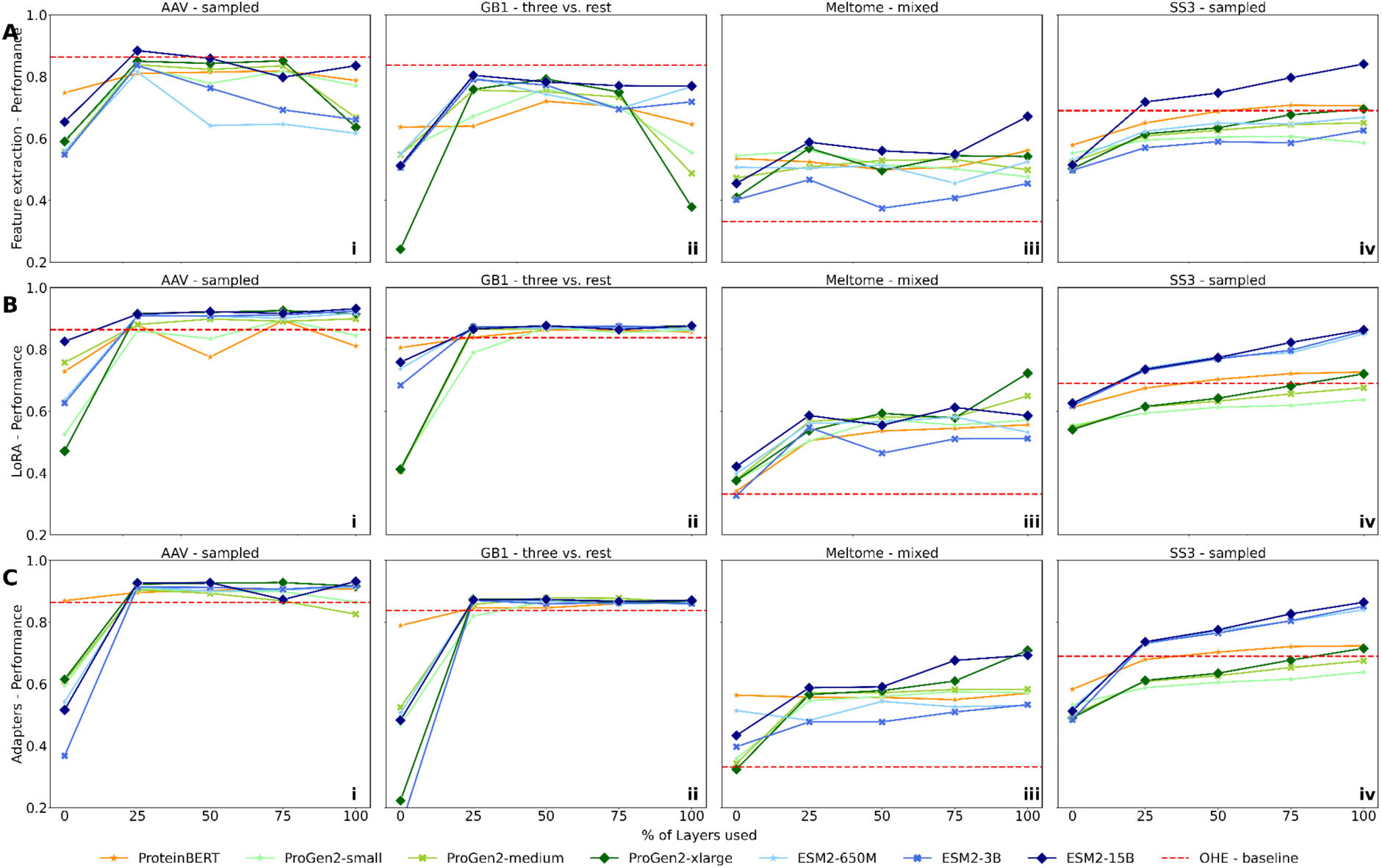
Performance analysis of TL models using different fractional layers of PLMs. TL is performed across three tasks, AAV-sampled, GB1-three vs. rest, Meltome-mixed, and SS3-sampled (columns i-iii) using variable number of PLM layers paired with TL strategies, Feature Extraction, LoRA and adapters (rows A-C). PLMs are differentiated by color, and performance curves are displayed for each task. For each subplot, x-axis shows the percentage of layers used - 0 (corresponds to using only the first layer of the model), 25%, 50%, 75%, 100% (Full model), and y-axis shows the Spearman’s correlation between the predicted and the ground truth values. The red dashed line represents a baseline model trained with the optimal hyperparameters using OHE of sequences, see Methods. PLM: Protein Language Models; TL: Transfer Learning; OHE: One Hot Encoding, FE: Feature Extraction; LoRA: Low Rank Adaptation.

### Fine-tuning yields substantial performance gain in complex tasks

To evaluate the effectiveness of TL for six fitness prediction tasks (*AAV-sampled, AAV-one vs. rest, GB1-three vs. rest, GB1-one vs. rest, Meltome-mixed, SS3-sampled*), we compared the performances of the best configurations (optimal layer and hyperparameters) for each technique on different PLMs. In addition to the original versions of LoRA and adapters, we included in our analysis two lightweight versions, namely LoRA- and adapters-, focusing on fine-tuning the output representations rather than the entire model. While the standard methods apply their fine-tuning modules across multiple or all layers of the model, the “-” versions restrict these modules to only the final layer of the PLM. Overall, most TL techniques yielded superior results compared to baseline models trained with OHE of sequences (Figure 4A). For the *AAV-sampled* and *GB1-three vs. rest* setups, the performance improvements were marginal, raising questions about the utility of PLMs and TL for these specific setups (Figure 4Ai, 4Aiii). These tasks are relatively simple because training data consists of sequences emerging from similar distributions (i.e., mutational variants of a WT) and are tested/validated on similarly structured data (Table 1).

**Figure 4.**
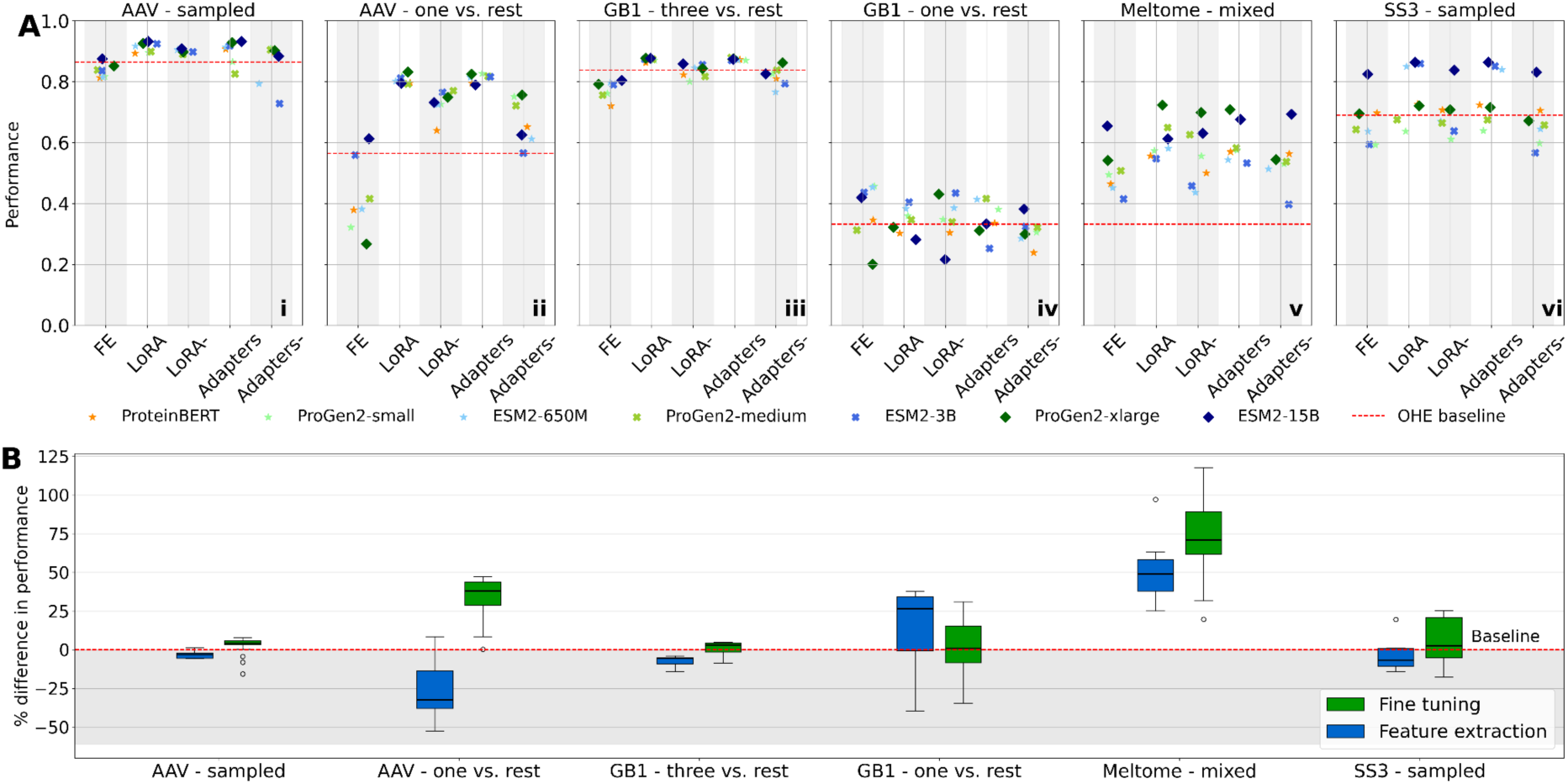
TL techniques performance comparison, in Spearman’s correlation (i-v), and macro accuracy (vi), across six different tasks. (A) Spearman’s correlation of the best performing PLM configuration with respect to the layer, downstream head and pooling method used for each TL technique (x-axis), is compared on: (i) *AAV-sampled*, (ii) *AAV-one vs. rest*, (iii) *GB1-three vs. rest*, (iv) *GB1-one vs. rest*, (v) *Meltome-mixed* and (vi) *SS3-sampled*. Different PLMs are used: ProteinBERT, ProGen2 (small, medium, xlarge), ESM2 (650M, 3B, 15B), with TL strategies including FE, LoRA, LoRA-, adapters, and adapters-. The red dashed line represents a baseline model trained with the optimal hyperparameters using OHE of sequences, see Methods. (B) Percentage difference in performance relative to baseline for FT (green) and FE (blue). Fine-tuning consistently yields larger performance improvements, particularly on more complex datasets like *Meltome - mixed*. Boxplots display variability in performance gains across tasks and TL methods. PLM: Protein Language Models; TL: Transfer Learning; OHE: One Hot Encoding; FE: Feature Extraction; FT: Fine-tuning; LoRA: Low Rank Adaptation

Conversely, TL methods demonstrated the highest performance gains for complex tasks consisting of training data with diverse protein sequences (*Meltome-mixed* and *SS3-sampled*), highlighting that PLMs excel in capturing distinct features across different protein families to accurately represent their amino acid sequences (Figure 4Av. 3vi). Similarly, TL methods also excelled in tasks where only single mutation variants were available during training, such as *AAV-one vs. rest* and *GB1-one vs. rest*, while sequences with higher ED were used for testing (Figure 4Aii, 3Aiv). This suggests that TL on PLMs enhances model generalization by leveraging pre-existing knowledge encoded in these models, which can be particularly advantageous when dealing with limited or single mutation datasets. Interestingly, for ProteinBERT and ProGen2 family of PLMs, where FE did not surpass the baseline model (Figure 4Aii), all FT methods appeared to recover and enhance performance, thus underscoring the importance of co-optimizing pre-trained PLM weights while incorporating task-specific knowledge. This effect is particularly evident in the *AAV one vs. rest* setup, where FE techniques averaged a Spearman’s correlation (*ρ*_⬚_) of 0.44, which is 21.5% below the baseline (*ρ* = 0.565), and whereas all FT models outperformed the baseline (average *ρ* = 0.75, +34%).

Figure 4B illustrates the distribution of FE and FT effects on performance (percentage changes) compared to baseline models across each of the six tasks and highlights their impact on task performance in relation to the complexity of the task. For the *AAV-sampled* and *GB1-three vs. rest* tasks, both FE and FT approaches perform similarly and close to the baseline with median performance difference of -3.2% / 3.95% and -5.64% / 2.89% respectively (Table S4). However, as task complexity increases, such as in the *Meltome-mixed* task, FT yields substantial performance gain compared to FE, which can achieve an improvement of up to 117.82% over the baseline methods (Table S4). Simultaneously optimizing novel parameters with pre-trained weights, FT allows the models to extract deeper representations from the PLMs, enabling more effective learning from diverse and complex protein sequences. These results suggest that complex tasks benefit significantly from pre-training, which leverages the full capacity of the PLMs to handle intricate sequence variability and relationships.

PLMs did not present substantial differences in performance despite differences in protein families and data scale (Table 2). This suggests that the PLM architecture and pretraining carries less significance than the TL technique choice and calibration. However, in the *Meltome-mixed* and *SS3-sampled* task we observe that the larger models (ProGen2-xlarge, ESM2-3B, ESM2-15B) produce superior performance, which is anticipated, as an increased number of parameters enhances the model’s capacity to represent the diversity of proteins in this task. These observations underscore that high performance can be attained by leveraging smaller models, provided that TL is applied effectively. The TL configuration that performed best for each task is presented in Table 2.

**Table 2.** Scoreboard of the best transfer learning configuration results for each task.

| Task | Score | Metric | Best configuration |  |  |  |  |
| --- | --- | --- | --- | --- | --- | --- | --- |
|  |  |  | PLM | TL-method | Layers used | Pooling | Downstream head |
| AAV one-vs-rest | 0.8315 | Spearman's Corr. | ProGen2-xlarge | LoRA | quarter3 | cls | linear |
| AAV sampled | 0.9320 | Spearman's Corr. | ESM2-15B | Adapters | last | mean | linear |
| GB1 one-vs-rest | 0.4571 | Spearman's Corr. | ProGen2-small | FE | quarter3 | mean | linear |
| GB1 three-vs-rest | 0.8791 | Spearman's Corr. | ProGen2-medium | Adapters | middle | cls | linear |
| Meltome mixed | 0.7232 | Spearman's Corr. | ProGen2-xlarge | LoRA | last | mean | linear |
| RBD one-vs-rest | 0.5545 | MCC | ProGen2-small | LoRA | middle | mean | linear |
| Trastuzumab one-vs-rest | 0.3898 | MCC | ProGen2-xlarge | LoRA- | quarter1 | cls | linear |
| SS3 sampled | 0.8644 | Macro Accuracy | ESM2-15B | Adapters | last | - | linear |

### Fine-tuning performs better to higher mutation variants when only labels for single mutations are available

Driven by the observation that fine-tuning PLMs can be particularly beneficial for *one vs. rest* splits, we determined the performance of the best configuration from each TL technique (Table 2) across varying degrees of ED when trained only with single mutation variants. All TL-based models maintained more consistent performance levels as the ED increases compared to the respective baseline model, which rapidly loses efficacy (Figure 5). Specifically for the AAV (Figure 5A) and RBD (Figure 5B) datasets, performance gradually declines at higher mutational levels, while the baseline models decline rapidly for the *AAV-one vs. rest*. Model training was not achievable for any OHE baseline model for the *RBD-one vs. rest* task (Figure 5B), likely due to the high sparsity introduced by OHE and/or the lack of capacity in logistic regression and single-layer neural networks architectures. LoRA- and adapter-based models trained for the *GB1-one vs. rest* and *Trastuzumab-one vs. rest* tasks outperformed FE and the baseline across all ED, except at ED = 4 for the GB1 dataset (Figure 5C, Figure 5D); however, this deviation in performance can likely be attributed to the small training data size in the GB1 dataset (Table 1), raising concerns regarding the reliability of the observed trends. Similarly, for the Trastuzumab dataset, the limited number of sequences at lower ED (Table 1) compromises the reliability of predictions at higher ED, rendering the results less conclusive.These observations imply that leveraging single mutation labeled data to guide PLMs via TL can yield models that combine general knowledge acquired during pre-training (i.e. fitness and evolution) with task-specific data (e.g. experimental), making them capable of effectively generalizing to previously unseen protein sequences.

**Figure 5.**
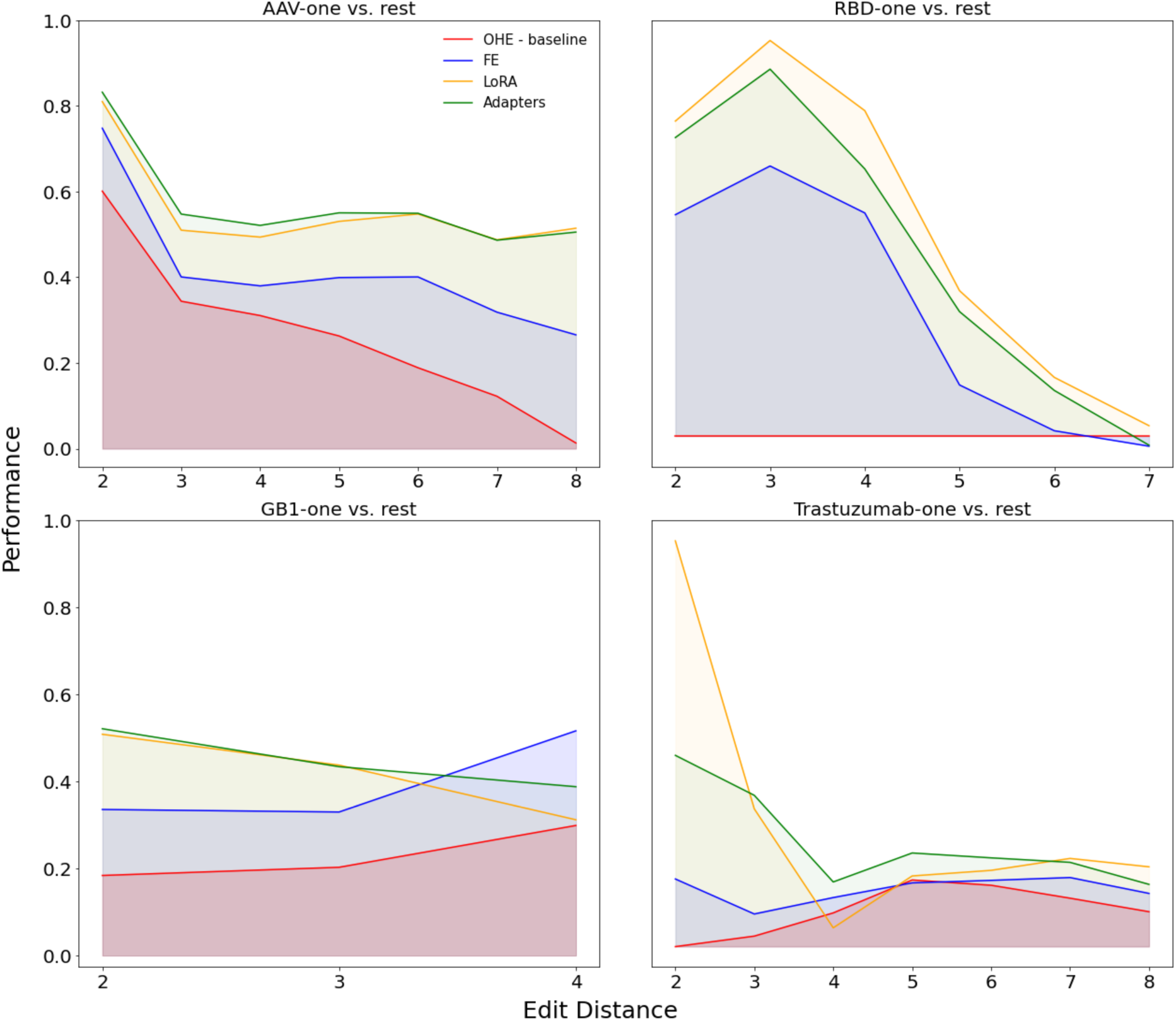
Performance of TL methods across different mutational edit distances for various datasets. (A-D) Performance trends as a function of edit distance for four different datasets: (A) AAV-one vs. rest, (B) RBD-one vs. rest, (C) GB1-one vs. rest, and (D) Trastuzumab-one vs. rest. The y-axis shows model performance (Spearman’s correlation for GB1 and AAV dataset, MCC for RBD and Trastuzumab dataset), while the x-axis represents the ED from the reference sequence. Each line represents a different transfer learning strategy. PLM: Protein Language Models; TL: Transfer Learning; OHE: One Hot Encoding; FE: Feature Extraction; FT: Fine-tuning; LoRA: Low Rank Adaptation; ED:Edit Distance; MCC: Matthews Correlation Coefficient

### Applying LoRA only on the last layer provides a reliable tradeoff between memory constraints and model performance

Our analysis demonstrated that FT techniques can yield superior results when adapting pre-trained PLMs for protein engineering tasks. Despite adopting PEFT methods, even when fine-tuning a small fraction of the model, the number of trainable parameters can still be large. This is due to the sheer size of models such as ESM2-15B and ProGen2-xlarge, which consist of 98 and 6.4 billion parameters, respectively. In such models, updating even a very small subset requires significant computational resources. Motivated by these challenges, we investigated the performance of FT modules added only to the final layer of the entire PLM, LoRA- and adapters-. Adopting this strategy, we reduced the number of trainable parameters by a fraction of the total layer number and reduced the computational and memory overhead. Comparative analysis reveals the performance (upper section) and memory usage (lower section) between LoRA, LoRA-, adapters, and adapters-across three tasks: *AAV-sampled*, *GB1-three vs. rest*, and *Meltome-mixe*d (Figure 6). While, both LoRA and adapters outperform their reduced counterparts, LoRA- and adapters-, the performance drops for *AAV-sampled* and *GB1-three vs rest* are marginal. Only *Meltome-mixed* exhibits a more notable decline when the adapters-method is applied. Importantly, both LoRA- and adapters- have significantly lower memory requirements, without a drastic performance loss. GPU RAM memory ranges from 6.5-12.8 gigabytes, compared to the standard application of these methods which require 28-52 gigabytes (Table 3). By reducing the memory requirements without significantly sacrificing performance, LoRA- and adapters- provide accessibility to larger PLMs and enable their FT without the need for expensive cloud computing services or specialized infrastructure. PLMs, training parameters and hardware resources used for LoRA- and adapters- for the comparative analysis and for the entirety of setups are shown in Table 3.

**Figure 6.**
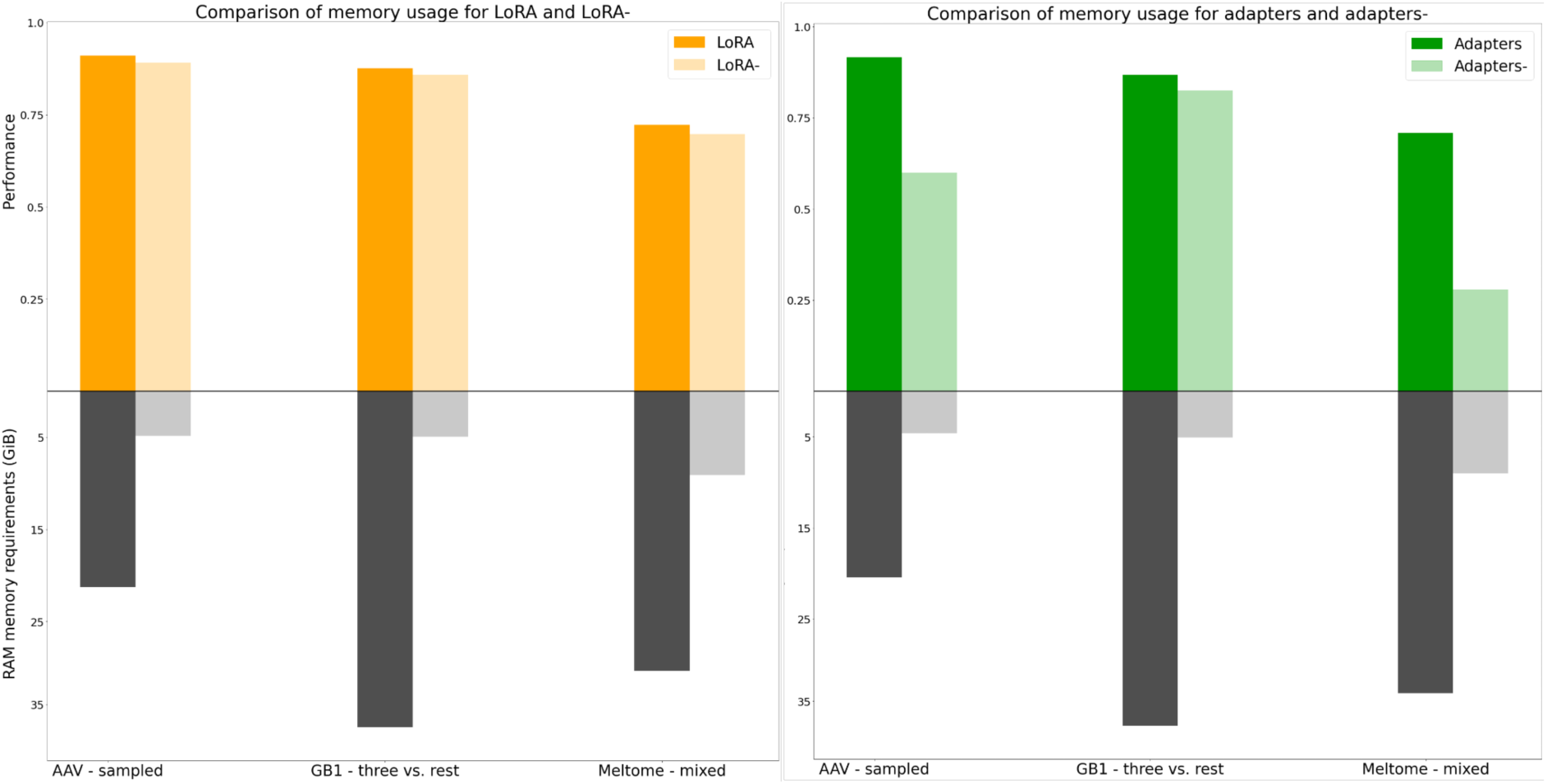
Performance and memory trade-off between TL techniques utilizing full or last layer only PLMs. The orange bars (dark for LoRA and light for LoRA-) and the green bars (dark for adapters and light for adapters-) show the performance (Spearman’s correlation) of the models on three datasets: *AAV-sampled, GB1-three vs. rest*, and *Meltome-mixed*. The y-axis represents performance on a scale from 0 to 1, where LoRA consistently outperforms LoRA-across all datasets. The gray part of the bars (dark for default and light for - version) represents the memory requirements in gigabytes (GB) for each approach. LoRA requires significantly more memory compared to LoRA-, illustrating a trade-off between performance and memory efficiency. LoRA: Low Rank Adaptation

**Table 3.** Model configuration for edit distance analysis.

| Tasks | PLM | Method | Performance | RAM GiB requirements | GPU Used |
| --- | --- | --- | --- | --- | --- |
| AAV - sampled | ESM-3B | LoRA | 0.911 | 28 | NVIDIA A100 80GB (x3) |
|  |  | LoRA- | 0.891 | 6.4 | NVIDIA Quadro RTX 6000 24G (x2) |
|  |  | Adapters | 0.917 | 29 | NVIDIA A100 80GB (x4) |
|  |  | Adapters- | 0.599 | 6.5 | NVIDIA Quadro RTX 6000 24G (x2) |
| GB1 - three vs. rest | ESM2-15B | LoRA | 0.876 | 48 | NVIDIA A100 80GB (x4) |
|  |  | LoRA- | 0.858 | 6.5 | NVIDIA Quadro RTX 6000 24G (x2) |
|  |  | Adapters | 0.868 | 52 | NVIDIA A100 80GB (x4) |
|  |  | Adapters- | 0.825 | 7.2 | NVIDIA Quadro RTX 6000 24G (x2) |
| Meltome - mixed | ProGen2-xlarge | LoRA | 0.723 | 40 | NVIDIA A100 80GB (x4) |
|  |  | LoRA- | 0.698 | 12 | NVIDIA Quadro RTX 6000 24G (x4) |
|  |  | Adapters | 0.708 | 47 | NVIDIA A100 80GB (x4) |
|  |  | Adapters- | 0.279 | 12.8 | NVIDIA Quadro RTX 6000 24G (x4) |

### Practical guidelines for applying PLMFit

Using the PLMfit platform, we have compiled practical guidelines for the research community to effectively apply TL techniques of FE and FT on PLMs. First, following the splitting of the dataset on training, validation, and testing sets, redundant (i.e., duplicates) and arbitrarily or mis-labeled (i.e., same amino acid sequence with different label) sequences must be removed to prevent ambiguity during training. Depending on the amount of diversity required for the task of interest, data can be further clustered by sequence identity with protein clustering tools (e.g., MMSeq2^32,33^ or kClust^32,33^). The main drivers of choosing a TL and PLM approach are the diversity of training data, the level of accuracy required for the task and amount of accessible resources. Generally, tasks performed on data with small sequence variation between the training and testing sets can benefit from training a custom model. However, choosing the optimal architecture and tuning the hyperparameters may not be straightforward, despite theoretically being able to bring higher results. In that case, FE using the 25% of a PLM with a linear downstream head could be sufficient and resource-efficient. Although fine-tuning a PLM can improve performance in these scenarios, the performance gains are often marginal.

For datasets characterized by single mutation variants (e.g., deep mutational scanning experiments), custom models typically struggle to generalize to variants with higher ED. In these instances, using LoRA to fine-tune a fraction (e.g. 25-75%) of a larger PLM (ESM2-15B, ProGen2-xlarge) is recommended to achieve better performance. Depending on the availability of GPUs, the LoRA-technique (i.e. fine-tuning only the last layer) can serve as an effective compromise, offering adequate performance yield while mitigating the computational burden for this task.

In more complex tasks involving diverse protein sequences (e.g., *Meltome-mixed*), model performance tends to scale with size and amount of trainable parameters. Consequently, employing TL on PLMs for this type of tasks can improve performance, as the extensive knowledge embedded in these models from large datasets of natural proteins can be effectively leveraged. Particularly, applying LoRA on large-scale PLMs, like ESM2-15B or ProGen2-xlarge, is likely to be the most effective method. However, when GPU availability or inference speed is a limiting factor, fine-tuning smaller PLMs, such as ESM-3B or ProGen2-medium, offers a practical alternative that can provide satisfactory performance.

Deepspeed package with CPU offloading and mixed precision training is utilized in PLMFit to manage computational resources^34^. Stage 3 of Deepspeed is applied, with smaller reduce and all-gather bucket sizes for resource-constrained setups. These values can be adjusted for faster processing. PyTorch Lightning is used for easier integration with Deepspeed, providing a streamlined setup and cleaner code. Fine-tuning a PLM on a specific dataset using PLMFit is a streamlined process that can be executed with a single command.

## DISCUSSION

In this study, we evaluated three TL approaches across six datasets, utilizing three families of state-of-the-art pre-trained PLMs. Each TL-based model was trained with varying parameters, including the number of layers used, pooling methods, downstream architecture, and training hyperparameters. Our analysis evaluates the effectiveness of transfer learning methods using protein language, extending prior research^23,27^ by exploring finer granularity in hyperparameter tuning for transfer learning and incorporating larger models. Performance was compared against task-specific baselines trained with OHE of sequences (Figure 4). The results indicate that when applied correctly, TL can offer substantial benefits for protein engineering-related tasks. Since protein language models are often used to describe sequences in an evolutionary context or guide evolutionary processes^8,35^, the insights from this work provide practical guidelines for leveraging stored knowledge effectively. FE can serve as valuable input features by incorporating evolutionary information into sequence encoding. FT, while improving performance across all tasks, may not always be the optimal choice due to marginal performance gains relative to the computational cost it incurs. In some cases, smaller models can achieve performance comparable to larger models when the optimal hyperparameters are applied for transfer learning, which can be particularly beneficial given the growing size and complexity of modern models^17^. Nevertheless, FT proves particularly advantageous in few-shot learning scenarios and more challenging tasks, enabling models to generalize to unseen data by leveraging pre-training information. In such scenarios, simultaneously optimizing novel parameters with pre-trained weights, FT allows the models to extract deeper representations from the PLMs, enabling more effective learning from diverse and complex protein sequences. A commonly used method in the protein engineering field is deep mutational scanning (DMS)^36^, which systematically introduces single mutations across a protein sequence and evaluates the resulting variants using an assay to measure a specific property of interest. DMS experiments are particularly valuable due to their high-throughput nature and relatively low experimental burden. Harnessing the information incorporated in DMS experiments to train or inform pre-trained PLMS limited data could be particularly valuable^25^ by enabling accurate predictions on combinatorial (i.e. ED > 1) libraries where multiple mutations have been introduced. Conversely, when a sufficient amount of data is available, and the predictive task involves variants with mutations at positions similar to those in the training data (i.e. the distribution between the training and test sets remains relatively close in terms of edit distance), the use of PLMs may be unnecessary, as a model trained ab-initio on the task-specific data could potentially offer comparable or even superior performance with greater efficiency. Thus, simpler, task-specific models might provide a more practical and effective solution for these types of problems, where the generalization capabilities of PLMs are less critical. Ultimately, the optimal use of pre-trained PLMs depends on the diversity and volume of training data, the specific nature of the task, and the computational resources available. As validated by earlier studies^22,28^, the last layer may not provide the optimal training features and in several setups even lead to substantial decrease in performance, this study reveals that targeting specific layers within the foundational model can not only enhance computational efficiency but also improve performance.

While our evaluation provides insights at a great scale and depth, a more robust evaluation scheme, such as k-fold cross-validation, could offer a more comprehensive assessment of each TL technique’s performance. Additionally, we did not explore an extensive space of ranks for LoRA and more complex architectures for adapters, due to the high computational and time costs associated with these approaches. In the future, a broader hyperparameter space should be investigated to effectively determine the impact of each parameter. Our comparative analysis focused on protein language models using only sequence data; however, the field is beginning to explore multimodal protein language models that integrate structural features^37,38^. The datasets and tasks used in this study can carry inherent biases, which could influence the results. To validate the initial observations from this study, it is essential to include a more diverse range of data taking into consideration a broad range of protein families and tasks. Ultimately, TL-based models will need to transition from computational validation to real-world lab experiments to truly demonstrate their potential. We also encourage the community to contribute by uploading new datasets, tasks, TL techniques, and additional pre-trained PLMs to help establish new benchmarks. Lastly, we intend to continuously update Table 2 with the highest performing combination of TL and PLM for different tasks to keep it relevant for ongoing research.

We anticipate that PLMFit will serve as a valuable resource for the research community by offering a practical starting point for those seeking to leverage PLMs for various protein engineering tasks. Whether users aim to fine-tune pre-trained PLMs or extract embeddings from different layers, PLMFit can act as a tool to streamline these processes. Additionally, the platform can guide researchers in selecting appropriate parameters and configurations, enabling the practical application of PLMs in a range of protein engineering tasks.

## METHODS

### Protein Language Models

TL techniques are applied to three state-of-the-art PLM families. Two BERT-based (ESM2 and ProteinBERT) and one GPT-based (ProGen2) foundational PLMs, pre-trained with MLM and CLM objectives respectively, are utilized. Different versions of these models are evaluated covering a broad range of architecture size with their layers size spanning from 12 to 48 layers, corresponding to 92M up to 15B pretrained parameters respectively. Analytic overview of the PLMs assessed in this study are shown in Table 4.

**Table 4.** Summary of protein language models used in this study.

| PLM | Type | No. of parameters | No. of layers | Embeddings Dim. |
| --- | --- | --- | --- | --- |
| ProteinBERT | MLM | 92M | 12 | 768 |
| ProGen2-small | CLM | 151M | 12 | 1024 |
| ProGen2-medium | CLM | 764M | 27 | 1536 |
| ProGen2-xlarge | CLM | 6.4B | 32 | 4096 |
| ESM2-650M | MLM | 650M | 33 | 1280 |
| ESM2-3B | MLM | 3B | 36 | 2560 |
| ESM2-15B | MLM | 15B | 48 | 5120 |

### Transfer Learning methods

Transfer learning entails reusing or adapting a pre-trained model as the foundation for addressing a different, novel task. By leveraging the knowledge from the original domain, performance on the new task can be improved. Depending on whether pre-trained weights (fully or partially) are co-optimized with the parameters of the newly added task specific downstream head, TL can be divided to feature Extraction and fine-tuning. To adequately examine if TL-based computational approaches can benefit protein engineering, we investigated both as part of this study.

#### Feature extraction

Feature extraction, as a TL method, involves the transformation of the input sequence into a numerical representation after performing inference on a pre-trained PLM without altering its weights. Conceptually, obtained representations encapsulate information about the structure and evolution of the processed protein sequence and can be used as input features to train shallow models in downstream tasks. In this study, we employ a layer pruning analysis assessing multiple fractions of the foundational models by extracting embeddings from the first, the last and three intermediate layers corresponding to 25%, 50%, and 75% of the models’ size.

#### Fine-tuning

On the contrary, during fine-tuning, weights of a pre-trained model are adjusted to the specific downstream task. Weight adaptation can occur either entirely or selectively by retraining a fraction of the model’s parameters while retaining the remaining frozen during back propagation. The latter approach significantly reduces the number of gradients to be calculated, thereby decreasing the time and resources required for the fine-tuning process. Parameter-Efficient Fine-Tuning (PEFT) extends this concept by combining novel trainable networks with the original parameters. By using the pre-trained model only for inference and updating solely the weights of the smaller, newly added, modules, knowledge acquired during pre-trained can be leveraged to optimize the model for the specific task of interest. Motivated by state-of-the-art techniques in Natural Language Processing and methods that have been studied in the biology realm, this study proposes two PLM Fine-tuning approaches to establish benchmarks, bottleneck adapters and Low-Rank Adaptation (LoRA). Adapters are small architectures injected between the layers of a pre-trained PLM, allowing for efficient fine-tuning by freezing the original model’s weights *W_o_* and only training the adapter parameters *W′* (1a). For this study, adapters’ architecture proposed by Yang et al.^24^ is employed. Low-Rank Adaptation (LoRA) decomposes the weight matrix *W_o_* of a pre-trained model into two low-rank (*r* ≪ *d_k_*) matrices *A* and *B*, significantly reducing the number of parameters to be trained (1b). LoRA modules applied on the pre-trained *W_q,k,v_* matrices of the attention heads in the different layers of the PLMs. Similarly, to feature extraction, the effect of adding FT-modules in different depths (first, last, intermediate; 25%, 50%, 75%) of PLMs is investigated. Additionally, to further decrease the amount of trainable parameters,we propose the addition of the respective modules only in the last layer of the foundation PLMs, namely LoRA- and the effect of this approach is being assessed as a trade-off between performance and computational efficiency.

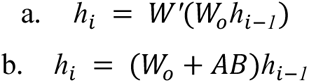

Equation (1). (a) Adapters fine-tuning where *h_i_* is the output of the current layer, *h_i-1_* is the input from the previous layer, *W_o_* is the weight matrix of the pre-trained model and *W′* is a small trainable matrix, referred to as the adapter, introduced to fine-tune the transformation for task-specific requirements. (b) Low-Rank Adaptation (LoRA) where *h_i_* is the output of the current layer, *h_i-1_* is the input from the previous layer, *W_o_* is the weight matrix of the pre-trained model, *A* and *B* are the trainable low-rank matrices which product *AB*modifies *W_o_* for fine-tuning with minimal computational overhead.

### Datasets and downstream tasks

Fitness prediction data acquired from the widely adopted Fitness Landscape Inference for Proteins (FLIP^15^) repository which includes curated datasets with experimental measurements that map protein sequences with binding affinity and thermostability measurements for different protein families. Particularly, it consists of two datasets of variants starting from a wild type sequence, Adeno-associated virus capsid (i.e. AAV) and GB1 domain of immunoglobulin-binding protein G (i.e. GB1), which fitness corresponds to variant enrichment ratio^39,40^ (i.e. E_r_). To simulate different training scenarios and use cases, each set is splitted in different setups. From the AAV dataset, we used “sampled” split, where sequences randomly chosen for training and testing, and “one vs rest” split, where single mutation variants used for training and the remaining sequences (up to 39 mutations) assigned to the evaluation set. From the GB1 dataset, “one vs rest” and “three vs rest” splits utilized for training the models with single and up to three edit distance variants respectively and the rest for testing (up to 4 mutations). For the thermostability prediction task, a dataset (i.e. Meltome) consisting of sequences clustered for 50% sequence similarity using MMSeq2^32^ have been assessed for the maximum temperature (i.e. T_o_) that they can perform their function (i.e. thermostability). A single split referred as “mixed” is used for this task, in which sequences randomly selected from 13 species are exploited during training.

The binding classification datasets used in this study consist of two distinct libraries^29,30^ screened for binding and escape against the human ACE2 receptor using yeast display and the HER2 antigen using mammalian display with hybridoma cells, respectively. The first library includes the mutational landscape of SARS-CoV-2 Omicron receptor-binding domain (RBD) variants, focusing on their binding and escape interactions with human ACE2. This library spans the entire 201 amino-acid RBD sequences, with mutations up to an edit distance of 7. The second library explores mutations within the complementarity-determining region H3 (CDRH3) of Trastuzumab, a therapeutic antibody, specifically assessing its binding affinity to the HER2 antigen. This dataset contains variants with up to 10 mutations in the Complementarity-determining regions (CDRH3) region.. We employ a similar splitting strategy as the FLIP repository for these datasets to define the binary classification tasks. For each, we define two splits: a “sampled” split, which consists of a random division of (70 % training, 15% validation, 15% testing), ensuring class balance in training and validation splits, and a “one-vs-rest” split, where only single-mutant variants (ED = 1) are used for training and validation, while variants with higher edit distances from the wild type (ED >1) are reserved for testing. Summary for all the data used in this study are shown in Table 1.

Finally, our analysis includes a secondary structure prediction (SS3) dataset for three-state secondary structure prediction (helix, strand, coil), derived from the nine-class DSSP 4.0^41^ annotations: α-helix (H), β-bridge (B), strand (E), 310-helix (G), π-helix (I), turn (T), bend (S), loop (L) and poly-proline helix (P). Following the procedure described by Yang et al.^24^, these nine labels are mapped to three states: H, G, I, P as helix, B, E as strand, and T, S, L as coil. This simpler three-state representation is common for secondary structure classification tasks, as it captures broad structural categories while reducing labeling noise. The final “sampled” split, also adopted from the source, comprises 9504 sequences for training, 1052 for validation, and 357 for testing, making SS3 a challenging task due to sequence diversity and the comprehensive coverage of different structural states.

### Evaluation metrics and baselines

In this study, the performance of each TL-based model was compared to hyperparameter-tuned (Table S2) logistic regression and multilayer perceptron (MLP) neural networks with one hidden layer. These comparisons were conducted using one-hot encodings (OHE). OHE is a method where each amino acid in a protein sequence is represented by a binary vector of length equal to the number of possible amino acids, where all elements are set to zero, except for the element in the position corresponding to the amino-acid in the protein sequence which is set to one. No information about the biochemical properties or evolutionary relationships between amino acids is captured using this encoding method. Following the encoding of each position with the amino acid corresponding one-hot vector, the input sequence represented by a 2-d matrix Z(*Z∈R^sequence length x embedding dimension^*) is flattened before being used as input features to train the baseline models.

The evaluation metrics implemented in this study vary based on the nature of the task. For models trained on regression tasks, we utilized Spearman’s correlation coefficient (*ρ*) to assess the strength and direction of the monotonic relationship between predicted and actual values (2).

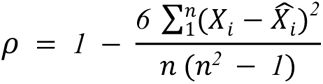

Equation (2). Spearman’s rank correlation coefficient (*ρ*) where *ρ∈*[−*1*,*1*], *X_i_* is the rank of the i-th element of real values, is the rank of the i-th element of predicted values, n is the total number of elements

For binary classification tasks, we employed Matthew’s correlation coefficient (MCC), which provides a balanced measure of the quality of binary classifications, taking into account true and false positives and negatives (3)

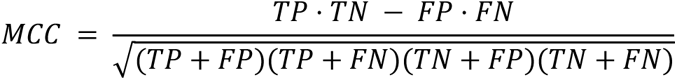

Equation (3). Matthew’s correlation coefficient (MCC) where TP, TN, FP, FN are the number of true positive, true negative, false positive and false negative prediction predictions respectively

For the secondary structure prediction task, where classification predictions are made per token (i.e., for each residue in the protein sequence), we evaluated the model using the macro accuracy of all tokens in a sequence, which provides an averaged measure of the model’s accuracy across all residues irrespective of class imbalance (4).

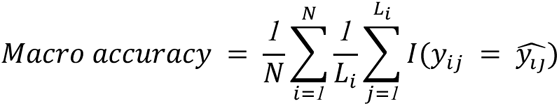

Equation 4. Macro accuracy, where *N* is the total number of protein sequences in the dataset, *L_i_* is the length of the i-th protein sequence, *y_ij_* and 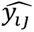 are the true and predicted labels for the j-th residue in the i-th sequence, respectively, and *I*(⋅) is an indicator function that equals 1 if the true label matches the predicted label and 0 otherwise.

### Training procedure

Each architecture evaluated in this study, whether based on transfer learning (TL) or initialized with random weights, is trained until convergence, with early stopping based on validation loss as the stopping criterion. For each FE-based setup, we trained the downstream modules using embeddings generated from PLMs (ESM, ProGen, and ProteinBERT). To convert local embeddings into a global representation, we explored a parameter space that included two pooling methods: averaging the embeddings or utilizing the CLS token. For the downstream classifier, we evaluated two potential architectures—a logistic regression model and a one-layer artificial neural network (ANN)—to determine the optimal approach. All combinations of these settings were subject to hyperparameter tuning to ensure optimal training performance. For FT, we limited the downstream architecture to logistic regression due to the computational burden associated with running these experiments. The hyperparameter tuning space for FT was more constrained, as the training process is time- and resources-intensive; the parameter space was therefore defined based on configurations from existing literature^24,42^. Detailed descriptions of the training procedures and hyperparameter tuning can be found in the supplementary materials (Table S2).

## ACKNOWLEDGMENTS

We would like to gratefully acknowledge ETH Zurich (Euler cluster) for providing computing resources and GPUS.

## AUTHOR CONTRIBUTIONS

T.B., E.S., and S.T.R.; investigation, T.B., E.S.; software, T.B., E.S.; supervision, S.T.R.; writing-original draft T.B., E.S., and S.T.R.

## DATA AND MATERIALS AVAILABILITY

All codes and datasets are available at https://github.com/LSSI-ETH/plmfit. All datasets can also be found in their original repo: AAV/GB1/Meltome (https://github.com/J-SNACKKB/FLIP), Trastuzumab (https://github.com/dahjan/DMS_opt), RBD (https://github.com/LSSI-ETH/Omicron_DML), and SS3 (https://github.com/fengtuan/LIFT_SS).

