## Supplementary Materials for "PLMFit : Benchmarking Transfer Learning with Protein Language Models for Protein Engineering"

### SUPPLEMENTARY ONLINE MATERIAL
**FOR**

### PLMFit : Benchmarking Transfer Learning with Protein Language Models for Protein Engineering

Thomas Bikias[1], Evangelos Stamkopoulos[1] and Sai. T. Reddy[1, 2]*

[1]Department of Biosystems Science and Engineering, ETH Zurich, Basel, Switzerland.

[2]Botnar Institute of Immune Engineering, Basel, Switzerland

#### A. Downstream heads architectures

Outputs from different PLMs' encoder layers are used as representations of protein sequence. The original decoder has been discarded and replaced with the task specific downstream head using these embeddings as input features for training. Transformer-based encoder outputs are 2-d matrices ($V_{local} \epsilon R^{sequence\ length\ x\ embedding\ dimension}$) where each residue (i.e, token) is described by a 1-d numerical vector ($V_{global} \epsilon R^{\ embedding\ dimension}$). Prior to inputting these representations into the downstream architecture, it is necessary to transition from local (i.e. token-wise) to global (i.e. sequence-wise) representations, thereby transforming the entire sequence into a feature vector. To achieve this, multiple reduction approaches can be applied. Within the scope of this study, two reduction techniques were assessed, mean- and sum- pooling. By respectively averaging or adding the elements towards the $sequence\ length$ dimension for each position through the embedding dimension 2-d matrices ( $V_{local}$) are transformed to 1-d vector ($V_{global}$).

Leveraging global representations of protein sequences, deep learning-based architectures are trained to address specific tasks. We evaluated two shallow architectures, logistic regression and a two-layer multilayer perceptron (MLP). Our focus was on highlighting the information encapsulated in the protein language models' (PLMs) embeddings , thus we utilized architectures that do not require extensive optimization as downstream models.

#### B. Training and hyperparameter tuning

Multiple (>3000) setups were assessed in this study. All models were developed using the PyTorch library. For every training procedure Adam optimizer is used with early stopping on best validation loss. All one-hot-encoding baselines and FE-based models have been tuned for optimal hyperparameters (i.e. learning rate, batch size, weight decay) using the Bayesian Optimization algorithm. FT implemented using the Deepspeed (Table S1) package and multiple hyperparameters have been assessed based on trial and error and existing literature. For LoRA, ranks of 4, 8, 16 and batch sizes of 2, 4, 6, 16, 32 were tested before concluding to the final hyperparameters. Similarly, for adapters, bottleneck dimensions of 16, 32, 64 and batch sizes of 4, 8, 16, 32 were examined before concluding to the ones used. All pre-trained PLMs downloaded either from HuggingFace or their original repo and adjusted to allow high-throughput Transfer Learning. All training hyperparameters used for each TL setup are shown in Table S2.

| Parameter | Value |
|---|---|
| Stage | 3 |
| Parameter Offload | Yes |
| Optimizer Offload | Yes |
| Offload Device | CPU |
| Sub Group Size | 1e12 |
| Overlap Comms. | Yes |
| Allgather Bucket Size | 2e8 |
| Reduce Bucket Size | 2e8 |

**Table S1. Deepspeed parameters used in PLMFit for fine-tuning setups.**

| | GB1 one-vs-rest | GB1 three-vs-rest | AAV one-vs-rest | AAV sampled | Meltome mixed | RBD one-vs-rest | Trastuzumab one-vs-rest | SS3 sampled |
|---|---|---|---|---|---|---|---|---|
| **Feature extraction / One hot encoding** | | | | | | | | |
| Hyperparameters Tuned | Learning rate, weight decay, batch size, hidden dimension (MLP) | | | | | | | |
| Learning rate space | 1e-2 - 1e-6 | | | | | | | |
| Weight decay space | 1e-1 - 1e-6 | | | | | | | |
| Batch size space | 8 - 128 | | | | | | | |
| Hidden dimension space (MLP) | 64 - 2048 | | | | | | | |
| Total trials Linear/MLP | 100/500 | | | | | | | |
| Initial random points | 20 | | | | | | | |
| Loss function | MSE | | | | | BCE | | Cross Entropy |
| Optimizer | Adam | | | | | | | |
| Epochs | 200 | | | | | | | |
| Patience (early stopping) | 30 | | | | | | | |
| **LoRA** | | | | | | | | |
| Modules Applied | Q, K, V | | | | | | | |
| LoRA rank | 8 | | | | | | | |
| LoRA alpha | 16 | | | | | | | |
| LoRA dropout | 0.1 | | | | | | | |
| Loss function | MSE | | | | | BCE | | Cross Entropy |
| Optimizer | Adam (DeepSpeedCPU) | | | | | | | |
| Batch size | 4 | | | | | | | |
| Epochs | 200 | 30 | 30 | 3 | 10 | 150 | 150 | 20 |
| Patience (early stopping) | 100 | 5 | - | 1 | 5 | - | - | 3 |
| Learning rate | 1e-4 | | | | | | | |
| Weight decay | 1e-2 | | | | | | | |
| **LoRA-** | | | | | | | | |
| Modules Applied | Q, K, V | | | | | | | |
| LoRA rank | 8 | | | | | | | |
| LoRA alpha | 16 | | | | | | | |
| LoRA dropout | 0.1 | | | | | | | |
| Loss function | MSE | | | | | BCE | | Cross Entropy |
| Optimizer | Adam (DeepSpeedCPU) | | | | | | | |
| Batch size | 4 | | | | | | | |
| Epochs | 200 | 30 | 30 | 10 | 15 | 150 | 150 | 30 |
| Patience (early stopping) | 100 | 5 | - | 5 | 5 | - | - | 3 |
| Learning rate | 1e-4 | | | | | | | |
| Weight decay | 1e-2 | | | | | | | |
| **Adapters** | | | | | | | | |
| Modules Applied | After FFN | | | | | | | |
| Bottleneck dimension | 32 | | | | | | | |
| Scaling | Learned | | | | | | | |
| Adapter dropout | 0.1 | | | | | | | |
| Loss function | MSE | | | | | BCE | | Cross Entropy |
| Optimizer | Adam (DeepSpeedCPU) | | | | | | | |
| Batch size | 4 | | | | | | | |
| Epochs | 200 | 30 | 30 | 3 | 10 | 150 | 150 | 20 |
| Patience (early stopping) | 100 | 5 | - | 1 | 5 | - | - | 3 |
| Learning rate | 1e-4 | | | | | | | |
| Weight decay | 1e-4 | | | | | | | |
| **Adapters-** | | | | | | | | |
| Modules Applied | After FFN | | | | | | | |
| Bottleneck dimension | 32 | | | | | | | |
| Scaling | Learned | | | | | | | |
| Adapter dropout | 0.1 | | | | | | | |
| Loss function | MSE | | | | | BCE | | Cross Entropy |
| Optimizer | Adam (DeepSpeedCPU) | | | | | | | |
| Batch size | 4 | | | | | | | |
| Epochs | 200 | 30 | 30 | 10 | 15 | 150 | 150 | 25 |
| Patience (early stopping) | 100 | 5 | - | 5 | 5 | - | - | 3 |
| Learning rate | 1e-4 | | | | | | | |
| Weight decay | 1e-4 | | | | | | | |

**Table S2. Hyperparameters used for all setups across the different tasks and methods.**

##### C. Hardware resources

For all experiments, we utilized ETH's high-performance computing cluster, Euler. The choice of hardware setup varied based on dataset size and transfer learning techniques. Multiple Nvidia GPUs (GeForce RTX 2080 Ti, RTX 3090, RTX 4090, TITAN RTX, Quadro RTX 6000, Tesla A100) were utilized for inference and backpropagation, with 1 to 4 GPUs used in parallel to accelerate training. Tables S3-S10 list the detailed resources used for each setup. Different pooling techniques require the same amount of resources and are therefore combined.

| PLM + Layer | Embeddings Extraction | | Feature Extraction | | LoRA | | LoRA- | | Adapters | | Adapters- | |
|---|---|---|---|---|---|---|---|---|---|---|---|---|
| | GPUs | CPU RAM | GPUs | CPU RAM | GPUs | CPU RAM | GPUs | CPU RAM | GPUs | CPU RAM | GPUs | CPU RAM |
| ProteinBERT - 1st | 1 x RTX 4090 | 12GB | 1 x RTX 4090 | 10GB | 1 x RTX 4090 | 12GB | 1 x RTX 2080 Ti | 12GB | 1 x RTX 4090 | 8GB | 1 x RTX 2080 Ti | 8GB |
| 25% | 1 x RTX 4090 | 12GB | 1 x RTX 4090 | 10GB | 1 x RTX 4090 | 12GB | 1 x RTX 2080 Ti | 12GB | 1 x RTX 4090 | 8GB | 1 x RTX 2080 Ti | 8GB |
| 50% | 1 x RTX 4090 | 12GB | 1 x RTX 4090 | 10GB | 1 x RTX 4090 | 12GB | 1 x RTX 2080 Ti | 12GB | 1 x RTX 4090 | 8GB | 1 x RTX 2080 Ti | 8GB |
| 75% | 1 x RTX 4090 | 12GB | 1 x RTX 4090 | 10GB | 1 x RTX 4090 | 12GB | 1 x RTX 2080 Ti | 12GB | 1 x RTX 4090 | 8GB | 1 x RTX 2080 Ti | 8GB |
| All/Last | 1 x RTX 4090 | 12GB | 1 x RTX 4090 | 10GB | 1 x RTX 4090 | 12GB | 1 x RTX 2080 Ti | 12GB | 1 x RTX 4090 | 8GB | 1 x RTX 2080 Ti | 8GB |
| ProGen2-small - 1st | 1 x RTX 4090 | 12GB | 1 x RTX 4090 | 10GB | 1 x RTX 4090 | 12GB | 1 x RTX 2080 Ti | 12GB | 1 x RTX 4090 | 12GB | 1 x RTX 2080 Ti | 12GB |
| 25% | 1 x RTX 4090 | 12GB | 1 x RTX 4090 | 10GB | 1 x RTX 4090 | 12GB | 1 x RTX 2080 Ti | 12GB | 1 x RTX 4090 | 12GB | 1 x RTX 2080 Ti | 12GB |
| 50% | 1 x RTX 4090 | 12GB | 1 x RTX 4090 | 10GB | 1 x RTX 4090 | 12GB | 1 x RTX 2080 Ti | 12GB | 1 x RTX 4090 | 12GB | 1 x RTX 2080 Ti | 12GB |
| 75% | 1 x RTX 4090 | 12GB | 1 x RTX 4090 | 10GB | 1 x RTX 4090 | 12GB | 1 x RTX 2080 Ti | 12GB | 1 x RTX 4090 | 12GB | 1 x RTX 2080 Ti | 12GB |
| All/Last | 1 x RTX 4090 | 12GB | 1 x RTX 4090 | 10GB | 1 x RTX 4090 | 12GB | 1 x RTX 2080 Ti | 12GB | 1 x RTX 4090 | 12GB | 1 x RTX 2080 Ti | 12GB |
| ESM2-650M - 1st | 1 x RTX 4090 | 12GB | 1 x RTX 4090 | 14GB | 1 x RTX 3090 | 12GB | 1 x RTX 2080 Ti | 12GB | 1 x RTX 4090 | 18GB | 1 x RTX 2080 Ti | 18GB |
| 25% | 1 x RTX 4090 | 12GB | 1 x RTX 4090 | 14GB | 1 x RTX 3090 | 12GB | 1 x RTX 2080 Ti | 12GB | 1 x RTX 4090 | 18GB | 1 x RTX 2080 Ti | 18GB |
| 50% | 1 x RTX 4090 | 12GB | 1 x RTX 4090 | 14GB | 2 x RTX 4090 | 12GB | 1 x RTX 2080 Ti | 12GB | 1 x RTX 4090 | 18GB | 1 x RTX 2080 Ti | 18GB |
| 75% | 1 x RTX 4090 | 12GB | 1 x RTX 4090 | 14GB | 2 x Quadro RTX 6000 | 12GB | 1 x RTX 2080 Ti | 12GB | 1 x RTX 4090 | 18GB | 1 x RTX 2080 Ti | 18GB |
| All/Last | 1 x RTX 4090 | 12GB | 1 x RTX 4090 | 14GB | 2 x Quadro RTX 6000 | 12GB | 2 x RTX 3090 | 12GB | 2 x RTX 3090 | 18GB | 2 x RTX 3090 | 18GB |
| ProGen2-medium - 1st | 1 x RTX 4090 | 12GB | 1 x RTX 4090 | 14GB | 1 x RTX 4090 | 12GB | 1 x RTX 2080 Ti | 12GB | 1 x RTX 4090 | 18GB | 1 x RTX 2080 Ti | 18GB |
| 25% | 1 x RTX 4090 | 12GB | 1 x RTX 4090 | 14GB | 1 x RTX 4090 | 12GB | 1 x RTX 2080 Ti | 12GB | 1 x RTX 4090 | 18GB | 1 x RTX 2080 Ti | 18GB |
| 50% | 1 x RTX 4090 | 12GB | 1 x RTX 4090 | 14GB | 2 x RTX 2080 Ti | 12GB | 1 x RTX 2080 Ti | 12GB | 1 x RTX 4090 | 18GB | 1 x RTX 2080 Ti | 18GB |
| 75% | 1 x RTX 4090 | 12GB | 1 x RTX 4090 | 14GB | 2 x TITAN RTX | 12GB | 1 x RTX 2080 Ti | 12GB | 1 x RTX 4090 | 18GB | 1 x RTX 2080 Ti | 18GB |
| All/Last | 1 x RTX 4090 | 12GB | 1 x RTX 4090 | 14GB | 2 x TITAN RTX | 12GB | 2 x RTX 3090 | 12GB | 2 x RTX 3090 | 18GB | 2 x RTX 3090 | 18GB |
| ESM2-3B - 1st | 1 x RTX 4090 | 30GB | 1 x RTX 4090 | 14GB | 1 x RTX 3090 | 40GB | 1 x RTX 2080 Ti | 40GB | 1 x RTX 4090 | 18GB | 1 x RTX 2080 Ti | 18GB |
| 25% | 1 x RTX 4090 | 30GB | 1 x RTX 4090 | 14GB | 1 x RTX 4090 | 40GB | 1 x RTX 2080 Ti | 40GB | 1 x RTX 4090 | 18GB | 1 x RTX 2080 Ti | 18GB |
| 50% | 1 x RTX 4090 | 30GB | 1 x RTX 4090 | 14GB | 1 x Quadro RTX 6000 | 40GB | 1 x RTX 2080 Ti | 40GB | 1 x RTX 4090 | 18GB | 1 x RTX 2080 Ti | 18GB |
| 75% | 1 x RTX 4090 | 30GB | 1 x RTX 4090 | 14GB | 1 x Quadro RTX 6000 | 40GB | 2 x RTX 3090 | 40GB | 2 x TITAN RTX | 18GB | 2 x RTX 3090 | 18GB |
| All/Last | 1 x RTX 4090 | 30GB | 1 x RTX 4090 | 14GB | 1 x A100 (40 GiB) | 40GB | 2 x TITAN RTX | 40GB | 2 x RTX 3090 | 18GB | 2 x TITAN RTX | 18GB |
| ProGen2-xlarge - 1st | 1 x RTX 4090 | 60GB | 1 x RTX 4090 | 14GB | 1 x RTX 2080 Ti | 40GB | 1 x RTX 2080 Ti | 40GB | 1 x RTX 2080 Ti | 40GB | 1 x RTX 2080 Ti | 40GB |
| 25% | 1 x RTX 4090 | 60GB | 1 x RTX 4090 | 14GB | 1 x RTX 2080 Ti | 40GB | 1 x RTX 2080 Ti | 40GB | 1 x RTX 2080 Ti | 40GB | 1 x RTX 2080 Ti | 40GB |
| 50% | 1 x RTX 4090 | 60GB | 1 x RTX 4090 | 14GB | 1 x RTX 2080 Ti | 40GB | 1 x RTX 2080 Ti | 40GB | 1 x RTX 3090 | 40GB | 1 x RTX 2080 Ti | 40GB |
| 75% | 1 x RTX 4090 | 60GB | 1 x RTX 4090 | 14GB | 1 x RTX 2080 Ti | 40GB | 2 x RTX 3090 | 40GB | 2 x TITAN RTX | 40GB | 2 x RTX 3090 | 40GB |
| All/Last | 1 x A100 (80 GiB) | 60GB | 1 x RTX 4090 | 14GB | 2 x TITAN RTX | 40GB | 2 x TITAN RTX | 40GB | 2 x RTX 3090 | 40GB | 2 x TITAN RTX | 40GB |
| ESM2-15B - 1st | 1 x RTX 4090 | 80GB | 1 x RTX 4090 | 14GB | 1 x RTX 3090 | 70GB | 1 x RTX 2080 Ti | 70GB | 1 x RTX 2080 Ti | 75GB | 1 x RTX 2080 Ti | 75GB |
| 25% | 1 x RTX 4090 | 80GB | 1 x RTX 4090 | 14GB | 1 x RTX 4090 | 70GB | 1 x RTX 2080 Ti | 70GB | 1 x RTX 2080 Ti | 75GB | 1 x RTX 2080 Ti | 75GB |
| 50% | 1 x RTX 4090 | 80GB | 1 x RTX 4090 | 14GB | 1 x Quadro RTX 6000 | 70GB | 1 x RTX 4090 | 70GB | 1 x RTX 3090 | 75GB | 2 x RTX 4090 | 75GB |
| 75% | 1 x A100 (80 GiB) | 80GB | 1 x RTX 4090 | 14GB | 1 x Quadro RTX 6000 | 70GB | 2 x TITAN RTX | 70GB | 2 x A100 (40 GiB) | 75GB | 2 x TITAN RTX | 75GB |
| All/Last | 1 x A100 (80 GiB) | 80GB | 1 x RTX 4090 | 14GB | 1 x A100 (40 GiB) | 70GB | 2 x A100 (40 GiB) | 70GB | 2 x A100 (80 GiB) | 75GB | 2 x A100 (80 GiB) | 75GB |

**Table S3. Detailed computational resources (number of GPUs and type, CPU RAM) used for each TL-based model on *GB1-one vs. rest* task.**

| PLM + Layer | Embeddings Extraction | | Feature Extraction | | LoRA | | LoRA- | | Adapters | | Adapters- | |
|---|---|---|---|---|---|---|---|---|---|---|---|---|
| | GPUs | CPU RAM | GPUs | CPU RAM | GPUs | CPU RAM | GPUs | CPU RAM | GPUs | CPU RAM | GPUs | CPU RAM |
| ProteinBERT - 1st | 1 x RTX 4090 | 12GB | 1 x RTX 4090 | 10GB | 1 x RTX 2080 Ti | 12GB | 1 x RTX 2080 Ti | 12GB | 1 x RTX 2080 Ti | 8GB | 1 x RTX 2080 Ti | 8GB |
| 25% | 1 x RTX 4090 | 12GB | 1 x RTX 4090 | 10GB | 1 x RTX 2080 Ti | 12GB | 1 x RTX 2080 Ti | 12GB | 1 x RTX 2080 Ti | 8GB | 1 x RTX 2080 Ti | 8GB |
| 50% | 1 x RTX 4090 | 12GB | 1 x RTX 4090 | 10GB | 1 x RTX 2080 Ti | 12GB | 1 x RTX 2080 Ti | 12GB | 1 x RTX 2080 Ti | 8GB | 1 x RTX 2080 Ti | 8GB |
| 75% | 1 x RTX 4090 | 12GB | 1 x RTX 4090 | 10GB | 1 x RTX 2080 Ti | 12GB | 1 x RTX 2080 Ti | 12GB | 1 x RTX 2080 Ti | 8GB | 1 x RTX 2080 Ti | 8GB |
| All/Last | 1 x RTX 4090 | 12GB | 1 x RTX 4090 | 10GB | 1 x RTX 2080 Ti | 12GB | 1 x RTX 2080 Ti | 12GB | 1 x RTX 2080 Ti | 8GB | 1 x RTX 2080 Ti | 8GB |
| ProGen2-small - 1st | 1 x RTX 4090 | 12GB | 1 x RTX 4090 | 10GB | 1 x RTX 2080 Ti | 12GB | 1 x RTX 2080 Ti | 12GB | 1 x RTX 2080 Ti | 12GB | 1 x RTX 2080 Ti | 12GB |
| 25% | 1 x RTX 4090 | 12GB | 1 x RTX 4090 | 10GB | 1 x RTX 2080 Ti | 12GB | 1 x RTX 2080 Ti | 12GB | 1 x RTX 2080 Ti | 12GB | 1 x RTX 2080 Ti | 12GB |
| 50% | 1 x RTX 4090 | 12GB | 1 x RTX 4090 | 10GB | 1 x RTX 2080 Ti | 12GB | 1 x RTX 2080 Ti | 12GB | 1 x RTX 2080 Ti | 12GB | 1 x RTX 2080 Ti | 12GB |
| 75% | 1 x RTX 4090 | 12GB | 1 x RTX 4090 | 10GB | 1 x RTX 2080 Ti | 12GB | 1 x RTX 2080 Ti | 12GB | 1 x RTX 2080 Ti | 12GB | 1 x RTX 2080 Ti | 12GB |
| All/Last | 1 x RTX 4090 | 12GB | 1 x RTX 4090 | 10GB | 1 x RTX 2080 Ti | 12GB | 1 x RTX 2080 Ti | 12GB | 1 x RTX 2080 Ti | 12GB | 1 x RTX 2080 Ti | 12GB |
| ESM2-650M - 1st | 1 x RTX 4090 | 12GB | 1 x RTX 4090 | 14GB | 1 x RTX 4090 | 12GB | 1 x RTX 3090 | 12GB | 1 x RTX 3090 | 18GB | 1 x RTX 3090 | 18GB |
| 25% | 1 x RTX 4090 | 12GB | 1 x RTX 4090 | 14GB | 1 x RTX 4090 | 12GB | 1 x RTX 3090 | 12GB | 1 x RTX 4090 | 18GB | 1 x RTX 3090 | 18GB |
| 50% | 1 x RTX 4090 | 12GB | 1 x RTX 4090 | 14GB | 2 x RTX 4090 | 12GB | 1 x RTX 4090 | 12GB | 2 x RTX 4090 | 18GB | 1 x RTX 4090 | 18GB |
| 75% | 1 x RTX 4090 | 12GB | 1 x RTX 4090 | 14GB | 2 x Quadro RTX 6000 | 12GB | 1 x RTX 4090 | 12GB | 2 x Quadro RTX 6000 | 18GB | 1 x RTX 4090 | 18GB |
| All/Last | 1 x RTX 4090 | 12GB | 1 x RTX 4090 | 14GB | 2 x Quadro RTX 6000 | 12GB | 1 x RTX 4090 | 12GB | 2 x Quadro RTX 6000 | 18GB | 1 x RTX 4090 | 18GB |
| ProGen2-medium - 1st | 1 x RTX 4090 | 12GB | 1 x RTX 4090 | 14GB | 1 x RTX 3090 | 12GB | 1 x RTX 3090 | 12GB | 1 x RTX 3090 | 18GB | 1 x RTX 3090 | 18GB |
| 25% | 1 x RTX 4090 | 12GB | 1 x RTX 4090 | 14GB | 1 x RTX 3090 | 12GB | 1 x RTX 3090 | 12GB | 1 x RTX 3090 | 18GB | 1 x RTX 3090 | 18GB |
| 50% | 1 x RTX 4090 | 12GB | 1 x RTX 4090 | 14GB | 2 x RTX 4090 | 12GB | 1 x RTX 4090 | 12GB | 2 x RTX 4090 | 18GB | 1 x RTX 4090 | 18GB |
| 75% | 1 x RTX 4090 | 12GB | 1 x RTX 4090 | 14GB | 2 x Quadro RTX 6000 | 12GB | 1 x RTX 4090 | 12GB | 2 x Quadro RTX 6000 | 18GB | 1 x RTX 4090 | 18GB |
| All/Last | 1 x RTX 4090 | 12GB | 1 x RTX 4090 | 14GB | 2 x Quadro RTX 6000 | 12GB | 1 x RTX 4090 | 12GB | 2 x Quadro RTX 6000 | 18GB | 1 x RTX 4090 | 18GB |
| ESM2-3B - 1st | 1 x RTX 4090 | 30GB | 1 x RTX 4090 | 14GB | 1 x RTX 4090 | 40GB | 1 x RTX 3090 | 40GB | 1 x RTX 3090 | 18GB | 1 x RTX 3090 | 18GB |
| 25% | 1 x RTX 4090 | 30GB | 1 x RTX 4090 | 14GB | 1 x RTX 4090 | 40GB | 1 x RTX 4090 | 40GB | 1 x RTX 4090 | 18GB | 1 x RTX 4090 | 18GB |
| 50% | 1 x RTX 4090 | 30GB | 1 x RTX 4090 | 14GB | 1 x Quadro RTX 6000 | 40GB | 1 x Quadro RTX 6000 | 40GB | 4 x Quadro RTX 6000 | 18GB | 2 x Quadro RTX 6000 | 18GB |
| 75% | 1 x RTX 4090 | 30GB | 1 x RTX 4090 | 14GB | 1 x Quadro RTX 6000 | 40GB | 1 x Quadro RTX 6000 | 40GB | 4 x Quadro RTX 6000 | 18GB | 2 x Quadro RTX 6000 | 18GB |
| All/Last | 1 x RTX 4090 | 30GB | 1 x RTX 4090 | 14GB | 1 x A100 (40 GiB) | 40GB | 2 x Quadro RTX 6000 | 40GB | 4 x Quadro RTX 6000 | 18GB | 2 x Quadro RTX 6000 | 18GB |
| ProGen2-xlarge - 1st | 1 x RTX 4090 | 60GB | 1 x RTX 4090 | 14GB | 1 x RTX 3090 | 40GB | 1 x RTX 3090 | 40GB | 1 x RTX 3090 | 40GB | 1 x RTX 3090 | 40GB |
| 25% | 1 x RTX 4090 | 60GB | 1 x RTX 4090 | 14GB | 1 x RTX 4090 | 40GB | 1 x RTX 4090 | 40GB | 1 x RTX 4090 | 40GB | 1 x RTX 4090 | 40GB |
| 50% | 1 x RTX 4090 | 60GB | 1 x RTX 4090 | 14GB | 1 x Quadro RTX 6000 | 40GB | 1 x Quadro RTX 6000 | 40GB | 4 x Quadro RTX 6000 | 40GB | 2 x Quadro RTX 6000 | 40GB |
| 75% | 1 x RTX 4090 | 60GB | 1 x RTX 4090 | 14GB | 1 x Quadro RTX 6000 | 40GB | 1 x Quadro RTX 6000 | 40GB | 4 x Quadro RTX 6000 | 40GB | 2 x Quadro RTX 6000 | 40GB |
| All/Last | 1 x A100 (80 GiB) | 60GB | 1 x RTX 4090 | 14GB | 1 x Quadro RTX 6000 | 40GB | 2 x Quadro RTX 6000 | 40GB | 4 x Quadro RTX 6000 | 40GB | 2 x Quadro RTX 6000 | 40GB |
| ESM2-15B - 1st | 1 x RTX 4090 | 80GB | 1 x RTX 4090 | 14GB | 1 x RTX 3090 | 70GB | 1 x RTX 3090 | 70GB | 1 x RTX 3090 | 75GB | 1 x RTX 3090 | 75GB |
| 25% | 1 x RTX 4090 | 80GB | 1 x RTX 4090 | 14GB | 2 x RTX 4090 | 70GB | 2 x RTX 4090 | 70GB | 2 x RTX 4090 | 75GB | 1 x RTX 4090 | 75GB |
| 50% | 1 x RTX 4090 | 80GB | 1 x RTX 4090 | 14GB | 3 x A100 (40 GiB) | 70GB | 2 x RTX 4090 | 70GB | 4 x A100 (40 GiB) | 75GB | 2 x RTX 4090 | 75GB |
| 75% | 1 x A100 (80 GiB) | 80GB | 1 x RTX 4090 | 14GB | 3 x A100 (80 GiB) | 70GB | 2 x Quadro RTX 6000 | 70GB | 4 x A100 (80 GiB) | 75GB | 2 x Quadro RTX 6000 | 75GB |
| All/Last | 1 x A100 (80 GiB) | 80GB | 1 x RTX 4090 | 14GB | 4 x A100 (80 GiB) | 70GB | 2 x Quadro RTX 6000 | 70GB | 4 x A100 (80 GiB) | 75GB | 2 x Quadro RTX 6000 | 75GB |

**Table S4. Detailed computational resources (number of GPUs and type, CPU RAM) used for each TL-based model on *GB1-three vs. rest* task.**

| PLM + Layer | Embeddings Extraction | | Feature Extraction | | LoRA | | LoRA- | | Adapters | | Adapters- | |
|---|---|---|---|---|---|---|---|---|---|---|---|---|
| | GPUs | CPU RAM | GPUs | CPU RAM | GPUs | CPU RAM | GPUs | CPU RAM | GPUs | CPU RAM | GPUs | CPU RAM |
| ProteinBERT - 1st | 1 x RTX 4090 | 12GB | 1 x RTX 4090 | 10GB | 1 x RTX 2080 Ti | 8GB | 1 x RTX 2080 Ti | 8GB | 1 x RTX 2080 Ti | 8GB | 1 x RTX 2080 Ti | 8GB |
| 25% | 1 x RTX 4090 | 12GB | 1 x RTX 4090 | 10GB | 1 x RTX 2080 Ti | 8GB | 1 x RTX 2080 Ti | 8GB | 1 x RTX 2080 Ti | 8GB | 1 x RTX 2080 Ti | 8GB |
| 50% | 1 x RTX 4090 | 12GB | 1 x RTX 4090 | 10GB | 1 x RTX 2080 Ti | 8GB | 1 x RTX 2080 Ti | 8GB | 1 x RTX 2080 Ti | 8GB | 1 x RTX 2080 Ti | 8GB |
| 75% | 1 x RTX 4090 | 12GB | 1 x RTX 4090 | 10GB | 1 x RTX 2080 Ti | 8GB | 1 x RTX 2080 Ti | 8GB | 1 x RTX 2080 Ti | 8GB | 1 x RTX 2080 Ti | 8GB |
| All/Last | 1 x RTX 4090 | 12GB | 1 x RTX 4090 | 10GB | 1 x RTX 2080 Ti | 8GB | 1 x RTX 2080 Ti | 8GB | 1 x RTX 2080 Ti | 8GB | 1 x RTX 2080 Ti | 8GB |
| ProGen2-small - 1st | 1 x RTX 4090 | 12GB | 1 x RTX 4090 | 10GB | 1 x RTX 2080 Ti | 12GB | 1 x RTX 2080 Ti | 12GB | 1 x RTX 2080 Ti | 12GB | 1 x RTX 2080 Ti | 12GB |
| 25% | 1 x RTX 4090 | 12GB | 1 x RTX 4090 | 10GB | 1 x RTX 2080 Ti | 12GB | 1 x RTX 2080 Ti | 12GB | 1 x RTX 2080 Ti | 12GB | 1 x RTX 2080 Ti | 12GB |
| 50% | 1 x RTX 4090 | 12GB | 1 x RTX 4090 | 10GB | 1 x RTX 2080 Ti | 12GB | 1 x RTX 2080 Ti | 12GB | 1 x RTX 2080 Ti | 12GB | 1 x RTX 2080 Ti | 12GB |
| 75% | 1 x RTX 4090 | 12GB | 1 x RTX 4090 | 10GB | 1 x RTX 2080 Ti | 12GB | 1 x RTX 2080 Ti | 12GB | 1 x RTX 2080 Ti | 12GB | 1 x RTX 2080 Ti | 12GB |
| All/Last | 1 x RTX 4090 | 12GB | 1 x RTX 4090 | 10GB | 1 x RTX 2080 Ti | 12GB | 1 x RTX 2080 Ti | 12GB | 1 x RTX 2080 Ti | 12GB | 1 x RTX 2080 Ti | 12GB |
| ESM2-650M - 1st | 1 x RTX 4090 | 12GB | 1 x RTX 4090 | 14GB | 1 x RTX 3090 | 18GB | 1 x RTX 3090 | 18GB | 1 x RTX 3090 | 18GB | 1 x RTX 3090 | 18GB |
| 25% | 1 x RTX 4090 | 12GB | 1 x RTX 4090 | 14GB | 1 x RTX 3090 | 18GB | 1 x RTX 3090 | 18GB | 1 x RTX 3090 | 18GB | 1 x RTX 3090 | 18GB |
| 50% | 1 x RTX 4090 | 12GB | 1 x RTX 4090 | 14GB | 2 x RTX 3090 | 18GB | 2 x RTX 4090 | 18GB | 2 x RTX 4090 | 18GB | 2 x RTX 4090 | 18GB |
| 75% | 1 x RTX 4090 | 12GB | 1 x RTX 4090 | 14GB | 2 x TITAN RTX | 18GB | 2 x Quadro RTX 6000 | 18GB | 2 x Quadro RTX 6000 | 18GB | 2 x Quadro RTX 6000 | 18GB |
| All/Last | 1 x RTX 4090 | 12GB | 1 x RTX 4090 | 14GB | 2 x TITAN RTX | 18GB | 2 x Quadro RTX 6000 | 18GB | 2 x RTX 4090 | 18GB | 2 x Quadro RTX 6000 | 18GB |
| ProGen2-medium - 1st | 1 x RTX 4090 | 12GB | 1 x RTX 4090 | 14GB | 1 x RTX 3090 | 18GB | 1 x RTX 2080 Ti | 18GB | 1 x RTX 3090 | 18GB | 1 x RTX 2080 Ti | 18GB |
| 25% | 1 x RTX 4090 | 12GB | 1 x RTX 4090 | 14GB | 1 x RTX 3090 | 18GB | 1 x RTX 2080 Ti | 18GB | 1 x RTX 3090 | 18GB | 1 x RTX 2080 Ti | 18GB |
| 50% | 1 x RTX 4090 | 12GB | 1 x RTX 4090 | 14GB | 2 x RTX 3090 | 18GB | 2 x RTX 2080 Ti | 18GB | 2 x RTX 3090 | 18GB | 2 x RTX 2080 Ti | 18GB |
| 75% | 1 x RTX 4090 | 12GB | 1 x RTX 4090 | 14GB | 2 x TITAN RTX | 18GB | 2 x RTX 4090 | 18GB | 2 x TITAN RTX | 18GB | 2 x RTX 4090 | 18GB |
| All/Last | 1 x RTX 4090 | 12GB | 1 x RTX 4090 | 14GB | 2 x TITAN RTX | 18GB | 2 x RTX 4090 | 18GB | 2 x TITAN RTX | 18GB | 2 x RTX 4090 | 18GB |
| ESM2-3B - 1st | 1 x RTX 4090 | 30GB | 1 x RTX 4090 | 14GB | 1 x RTX 3090 | 18GB | 1 x RTX 3090 | 18GB | 1 x RTX 2080 Ti | 18GB | 1 x RTX 3090 | 18GB |
| 25% | 1 x RTX 4090 | 30GB | 1 x RTX 4090 | 14GB | 1 x RTX 3090 | 18GB | 1 x RTX 4090 | 18GB | 1 x RTX 3090 | 18GB | 1 x RTX 4090 | 18GB |
| 50% | 1 x RTX 4090 | 30GB | 1 x RTX 4090 | 14GB | 2 x RTX 3090 | 18GB | 1 x Quadro RTX 6000 | 18GB | 1 x RTX 3090 | 18GB | 1 x Quadro RTX 6000 | 18GB |
| 75% | 1 x RTX 4090 | 30GB | 1 x RTX 4090 | 14GB | 2 x TITAN RTX | 18GB | 1 x Quadro RTX 6000 | 18GB | 2 x A100 (40 GiB) | 18GB | 1 x Quadro RTX 6000 | 18GB |
| All/Last | 1 x RTX 4090 | 30GB | 1 x RTX 4090 | 14GB | 2 x TITAN RTX | 18GB | 1 x A100 (40 GiB) | 18GB | 3 x A100 (40 GiB) | 18GB | 1 x A100 (40 GiB) | 18GB |
| ProGen2-xlarge - 1st | 1 x RTX 4090 | 60GB | 1 x RTX 4090 | 14GB | 1 x RTX 2080 Ti | 40GB | 1 x RTX 2080 Ti | 40GB | 1 x RTX 2080 Ti | 40GB | 1 x RTX 2080 Ti | 40GB |
| 25% | 1 x RTX 4090 | 60GB | 1 x RTX 4090 | 14GB | 1 x RTX 3090 | 40GB | 1 x RTX 3090 | 40GB | 1 x RTX 3090 | 40GB | 1 x RTX 3090 | 40GB |
| 50% | 1 x RTX 4090 | 60GB | 1 x RTX 4090 | 14GB | 2 x RTX 3090 | 40GB | 2 x RTX 4090 | 40GB | 1 x RTX 3090 | 40GB | 2 x RTX 4090 | 40GB |
| 75% | 1 x RTX 4090 | 60GB | 1 x RTX 4090 | 14GB | 2 x A100 (40 GiB) | 40GB | 2 x RTX 4090 | 40GB | 2 x A100 (40 GiB) | 40GB | 2 x RTX 4090 | 40GB |
| All/Last | 1 x A100 (80 GiB) | 60GB | 1 x RTX 4090 | 14GB | 2 x A100 (80 GiB) | 40GB | 3 x TITAN RTX | 40GB | 3 x A100 (40 GiB) | 40GB | 3 x TITAN RTX | 40GB |
| ESM2-15B - 1st | 1 x RTX 4090 | 80GB | 1 x RTX 4090 | 14GB | 1 x RTX 2080 Ti | 70GB | 1 x RTX 3090 | 70GB | 1 x RTX 2080 Ti | 75GB | 1 x RTX 3090 | 75GB |
| 25% | 1 x RTX 4090 | 80GB | 1 x RTX 4090 | 14GB | 1 x RTX 3090 | 70GB | 1 x RTX 4090 | 70GB | 2 x RTX 3090 | 75GB | 2 x RTX 4090 | 75GB |
| 50% | 1 x RTX 4090 | 80GB | 1 x RTX 4090 | 14GB | 2 x A100 (40 GiB) | 70GB | 2 x Quadro RTX 6000 | 70GB | 3 x A100 (40 GiB) | 75GB | 2 x Quadro RTX 6000 | 75GB |
| 75% | 1 x A100 (80 GiB) | 80GB | 1 x RTX 4090 | 14GB | 3 x A100 (80 GiB) | 70GB | 2 x Quadro RTX 6000 | 70GB | 3 x A100 (80 GiB) | 75GB | 2 x Quadro RTX 6000 | 75GB |
| All/Last | 1 x A100 (80 GiB) | 80GB | 1 x RTX 4090 | 14GB | 3 x A100 (80 GiB) | 70GB | 2 x A100 (40 GiB) | 70GB | 4 x A100 (80 GiB) | 75GB | 2 x A100 (80 GiB) | 75GB |

**Table S5. Detailed computational resources (number of GPUs and type, CPU RAM) used for each TL-based model on *AAV-one vs. rest* task.**

| PLM + Layer | Embeddings Extraction | | Feature Extraction | | LoRA | | LoRA- | | Adapters | | Adapters- | |
|---|---|---|---|---|---|---|---|---|---|---|---|---|
| | GPUs | CPU RAM | GPUs | CPU RAM | GPUs | CPU RAM | GPUs | CPU RAM | GPUs | CPU RAM | GPUs | CPU RAM |
| ProteinBERT - 1st | 1 x RTX 4090 | 12GB | 1 x RTX 4090 | 10GB | 2 x RTX 2080 Ti | 12GB | 1 x RTX 2080 Ti | 12GB | 2 x RTX 2080 Ti | 8GB | 1 x RTX 2080 Ti | 8GB |
| 25% | 1 x RTX 4090 | 12GB | 1 x RTX 4090 | 10GB | 2 x RTX 2080 Ti | 12GB | 1 x RTX 2080 Ti | 12GB | 2 x RTX 2080 Ti | 8GB | 1 x RTX 2080 Ti | 8GB |
| 50% | 1 x RTX 4090 | 12GB | 1 x RTX 4090 | 10GB | 3 x RTX 2080 Ti | 12GB | 1 x RTX 2080 Ti | 12GB | 3 x RTX 2080 Ti | 8GB | 1 x RTX 2080 Ti | 8GB |
| 75% | 1 x RTX 4090 | 12GB | 1 x RTX 4090 | 10GB | 3 x RTX 2080 Ti | 12GB | 1 x RTX 2080 Ti | 12GB | 3 x RTX 2080 Ti | 8GB | 1 x RTX 2080 Ti | 8GB |
| All/Last | 1 x RTX 4090 | 12GB | 1 x RTX 4090 | 10GB | 3 x RTX 2080 Ti | 12GB | 1 x RTX 2080 Ti | 12GB | 3 x RTX 2080 Ti | 8GB | 1 x RTX 2080 Ti | 8GB |
| ProGen2-small - 1st | 1 x RTX 4090 | 12GB | 1 x RTX 4090 | 10GB | 2 x RTX 2080 Ti | 12GB | 1 x RTX 2080 Ti | 12GB | 2 x RTX 2080 Ti | 12GB | 1 x RTX 2080 Ti | 12GB |
| 25% | 1 x RTX 4090 | 12GB | 1 x RTX 4090 | 10GB | 2 x RTX 2080 Ti | 12GB | 1 x RTX 2080 Ti | 12GB | 2 x RTX 2080 Ti | 12GB | 1 x RTX 2080 Ti | 12GB |
| 50% | 1 x RTX 4090 | 12GB | 1 x RTX 4090 | 10GB | 3 x RTX 2080 Ti | 12GB | 1 x RTX 2080 Ti | 12GB | 3 x RTX 2080 Ti | 12GB | 1 x RTX 2080 Ti | 12GB |
| 75% | 1 x RTX 4090 | 12GB | 1 x RTX 4090 | 10GB | 3 x RTX 2080 Ti | 12GB | 1 x RTX 2080 Ti | 12GB | 3 x RTX 2080 Ti | 12GB | 1 x RTX 2080 Ti | 12GB |
| All/Last | 1 x RTX 4090 | 12GB | 1 x RTX 4090 | 10GB | 3 x RTX 2080 Ti | 12GB | 1 x RTX 2080 Ti | 12GB | 3 x RTX 2080 Ti | 12GB | 1 x RTX 2080 Ti | 12GB |
| ESM2-650M - 1st | 1 x RTX 4090 | 12GB | 1 x RTX 4090 | 14GB | 2 x RTX 4090 | 12GB | 1 x RTX 3090 | 12GB | 2 x RTX 4090 | 18GB | 1 x RTX 3090 | 18GB |
| 25% | 1 x RTX 4090 | 12GB | 1 x RTX 4090 | 14GB | 2 x RTX 4090 | 12GB | 1 x RTX 3090 | 12GB | 2 x RTX 4090 | 18GB | 1 x RTX 3090 | 18GB |
| 50% | 1 x RTX 4090 | 12GB | 1 x RTX 4090 | 14GB | 3 x RTX 4090 | 12GB | 2 x RTX 4090 | 12GB | 3 x RTX 4090 | 18GB | 2 x RTX 4090 | 18GB |
| 75% | 1 x RTX 4090 | 12GB | 1 x RTX 4090 | 14GB | 4 x RTX 4090 | 12GB | 2 x Quadro RTX 6000 | 12GB | 4 x RTX 4090 | 18GB | 2 x Quadro RTX 6000 | 18GB |
| All/Last | 1 x RTX 4090 | 12GB | 1 x RTX 4090 | 14GB | 4 x Quadro RTX 6000 | 12GB | 2 x Quadro RTX 6000 | 12GB | 4 x Quadro RTX 6000 | 18GB | 2 x Quadro RTX 6000 | 18GB |
| ProGen2-medium - 1st | 1 x RTX 4090 | 12GB | 1 x RTX 4090 | 14GB | 2 x RTX 4090 | 12GB | 1 x RTX 3090 | 12GB | 2 x RTX 4090 | 18GB | 1 x RTX 3090 | 18GB |
| 25% | 1 x RTX 4090 | 12GB | 1 x RTX 4090 | 14GB | 2 x RTX 4090 | 12GB | 1 x RTX 3090 | 12GB | 2 x RTX 4090 | 18GB | 1 x RTX 3090 | 18GB |
| 50% | 1 x RTX 4090 | 12GB | 1 x RTX 4090 | 14GB | 3 x RTX 4090 | 12GB | 2 x RTX 4090 | 12GB | 3 x RTX 4090 | 18GB | 2 x RTX 4090 | 18GB |
| 75% | 1 x RTX 4090 | 12GB | 1 x RTX 4090 | 14GB | 4 x RTX 4090 | 12GB | 2 x RTX 4090 | 12GB | 4 x RTX 4090 | 18GB | 2 x RTX 4090 | 18GB |
| All/Last | 1 x RTX 4090 | 12GB | 1 x RTX 4090 | 14GB | 4 x Quadro RTX 6000 | 12GB | 2 x RTX 4090 | 12GB | 4 x Quadro RTX 6000 | 18GB | 2 x RTX 4090 | 18GB |
| ESM2-3B - 1st | 1 x RTX 4090 | 30GB | 1 x RTX 4090 | 14GB | 2 x TITAN RTX | 40GB | 2 x RTX 3090 | 40GB | 2 x TITAN RTX | 18GB | 2 x RTX 3090 | 18GB |
| 25% | 1 x RTX 4090 | 30GB | 1 x RTX 4090 | 14GB | 2 x TITAN RTX | 40GB | 3 x RTX 4090 | 40GB | 2 x TITAN RTX | 18GB | 3 x RTX 4090 | 18GB |
| 50% | 1 x RTX 4090 | 30GB | 1 x RTX 4090 | 14GB | 3 x RTX 4090 | 40GB | 4 x TITAN RTX | 40GB | 4 x RTX 4090 | 18GB | 4 x TITAN RTX | 18GB |
| 75% | 1 x RTX 4090 | 30GB | 1 x RTX 4090 | 14GB | 3 x A100 (40 GiB) | 40GB | 4 x TITAN RTX | 40GB | 3 x A100 (40 GiB) | 18GB | 4 x TITAN RTX | 18GB |
| All/Last | 1 x RTX 4090 | 30GB | 1 x RTX 4090 | 14GB | 4 x A100 (40 GiB) | 40GB | 4 x Quadro RTX 6000 | 40GB | 4 x A100 (80 GiB) | 18GB | 4 x Quadro RTX 6000 | 18GB |
| ProGen2-xlarge - 1st | 1 x RTX 4090 | 60GB | 1 x RTX 4090 | 14GB | 2 x TITAN RTX | 40GB | 2 x RTX 3090 | 40GB | 2 x TITAN RTX | 40GB | 2 x RTX 3090 | 40GB |
| 25% | 1 x RTX 4090 | 60GB | 1 x RTX 4090 | 14GB | 2 x TITAN RTX | 40GB | 3 x RTX 4090 | 40GB | 2 x TITAN RTX | 40GB | 3 x RTX 4090 | 40GB |
| 50% | 1 x RTX 4090 | 60GB | 1 x RTX 4090 | 14GB | 4 x RTX 4090 | 40GB | 4 x TITAN RTX | 40GB | 4 x RTX 4090 | 40GB | 4 x TITAN RTX | 40GB |
| 75% | 1 x RTX 4090 | 60GB | 1 x RTX 4090 | 14GB | 4 x A100 (40 GiB) | 40GB | 4 x TITAN RTX | 40GB | 4 x A100 (40 GiB) | 40GB | 4 x TITAN RTX | 40GB |
| All/Last | 1 x A100 (80 GiB) | 60GB | 1 x RTX 4090 | 14GB | 4 x A100 (40 GiB) | 40GB | 4 x Quadro RTX 6000 | 40GB | 4 x A100 (40 GiB) | 40GB | 4 x Quadro RTX 6000 | 40GB |
| ESM2-15B - 1st | 1 x RTX 4090 | 80GB | 1 x RTX 4090 | 14GB | 2 x TITAN RTX | 70GB | 1 x RTX 3090 | 70GB | 2 x TITAN RTX | 75GB | 2 x RTX 3090 | 75GB |
| 25% | 1 x RTX 4090 | 80GB | 1 x RTX 4090 | 14GB | 3 x TITAN RTX | 70GB | 2 x Quadro RTX 6000 | 70GB | 3 x TITAN RTX | 75GB | 2 x TITAN RTX | 75GB |
| 50% | 1 x RTX 4090 | 80GB | 1 x RTX 4090 | 14GB | 3 x A100 (40 GiB) | 70GB | 3 x Quadro RTX 6000 | 70GB | 3 x A100 (40 GiB) | 75GB | 3 x TITAN RTX | 75GB |
| 75% | 1 x A100 (80 GiB) | 80GB | 1 x RTX 4090 | 14GB | 3 x A100 (80 GiB) | 70GB | 4 x Quadro RTX 6000 | 70GB | 4 x A100 (80 GiB) | 75GB | 4 x A100 (80 GiB) | 75GB |
| All/Last | 1 x A100 (80 GiB) | 80GB | 1 x RTX 4090 | 14GB | 4 x A100 (80 GiB) | 70GB | 4 x A100 (80 GiB) | 70GB | 4 x A100 (80 GiB) | 75GB | 4 x Quadro RTX 6000 | 75GB |

**Table S6. Detailed computational resources (number of GPUs and type, CPU RAM) used for each TL-based model on *AAV-sampled* task.**

| PLM + Layer | Embeddings Extraction | | Feature Extraction | | LoRA | | LoRA- | | Adapters | | Adapters- | |
|---|---|---|---|---|---|---|---|---|---|---|---|---|
| | GPUs | CPU RAM | GPUs | CPU RAM | GPUs | CPU RAM | GPUs | CPU RAM | GPUs | CPU RAM | GPUs | CPU RAM |
| ProteinBERT - 1st | 1 x RTX 4090 | 12GB | 1 x RTX 4090 | 10GB | 2 x RTX 2080 Ti | 12GB | 2 x RTX 3090 | 12GB | 2 x RTX 3090 | 8GB | 2 x RTX 3090 | 8GB |
| 25% | 1 x RTX 4090 | 12GB | 1 x RTX 4090 | 10GB | 2 x RTX 2080 Ti | 12GB | 2 x RTX 3090 | 12GB | 2 x RTX 3090 | 8GB | 2 x RTX 3090 | 8GB |
| 50% | 1 x RTX 4090 | 12GB | 1 x RTX 4090 | 10GB | 3 x RTX 2080 Ti | 12GB | 3 x RTX 2080 Ti | 12GB | 3 x RTX 4090 | 8GB | 3 x RTX 2080 Ti | 8GB |
| 75% | 1 x RTX 4090 | 12GB | 1 x RTX 4090 | 10GB | 3 x RTX 3090 | 12GB | 3 x RTX 3090 | 12GB | 3 x RTX 4090 | 8GB | 3 x RTX 3090 | 8GB |
| All/Last | 1 x RTX 4090 | 12GB | 1 x RTX 4090 | 10GB | 3 x RTX 3090 | 12GB | 3 x RTX 3090 | 12GB | 3 x Quadro RTX 6000 | 8GB | 3 x RTX 3090 | 8GB |
| ProGen2-small - 1st | 1 x RTX 4090 | 12GB | 1 x RTX 4090 | 10GB | 2 x RTX 2080 Ti | 12GB | 2 x RTX 2080 Ti | 12GB | 2 x RTX 3090 | 12GB | 2 x RTX 2080 Ti | 12GB |
| 25% | 1 x RTX 4090 | 12GB | 1 x RTX 4090 | 10GB | 2 x RTX 2080 Ti | 12GB | 2 x RTX 2080 Ti | 12GB | 2 x RTX 3090 | 12GB | 2 x RTX 2080 Ti | 12GB |
| 50% | 1 x RTX 4090 | 12GB | 1 x RTX 4090 | 10GB | 3 x RTX 2080 Ti | 12GB | 3 x RTX 2080 Ti | 12GB | 3 x TITAN RTX | 12GB | 3 x RTX 2080 Ti | 12GB |
| 75% | 1 x RTX 4090 | 12GB | 1 x RTX 4090 | 10GB | 3 x RTX 3090 | 12GB | 3 x RTX 3090 | 12GB | 3 x RTX 4090 | 12GB | 3 x RTX 3090 | 12GB |
| All/Last | 1 x RTX 4090 | 12GB | 1 x RTX 4090 | 10GB | 3 x RTX 3090 | 12GB | 3 x RTX 3090 | 12GB | 3 x RTX 4090 | 12GB | 3 x RTX 3090 | 12GB |
| ESM2-650M - 1st | 1 x RTX 4090 | 12GB | 1 x RTX 4090 | 14GB | 1 x RTX 4090 | 12GB | 1 x RTX 4090 | 12GB | 2 x RTX 4090 | 18GB | 1 x RTX 4090 | 18GB |
| 25% | 1 x RTX 4090 | 12GB | 1 x RTX 4090 | 14GB | 1 x RTX 4090 | 12GB | 1 x RTX 4090 | 12GB | 2 x RTX 4090 | 18GB | 1 x RTX 4090 | 18GB |
| 50% | 1 x RTX 4090 | 12GB | 1 x RTX 4090 | 14GB | 2 x RTX 4090 | 12GB | 2 x RTX 4090 | 12GB | 3 x RTX 4090 | 18GB | 2 x RTX 4090 | 18GB |
| 75% | 1 x RTX 4090 | 12GB | 1 x RTX 4090 | 14GB | 2 x RTX 4090 | 12GB | 2 x RTX 4090 | 12GB | 4 x RTX 4090 | 18GB | 2 x RTX 4090 | 18GB |
| All/Last | 1 x RTX 4090 | 12GB | 1 x RTX 4090 | 14GB | 4 x Quadro RTX 6000 | 12GB | 2 x TITAN RTX | 12GB | 4 x Quadro RTX 6000 | 18GB | 2 x TITAN RTX | 18GB |
| ProGen2-medium - 1st | 1 x RTX 4090 | 12GB | 1 x RTX 4090 | 14GB | 1 x RTX 4090 | 12GB | 1 x RTX 4090 | 12GB | 2 x RTX 4090 | 18GB | 1 x RTX 4090 | 18GB |
| 25% | 1 x RTX 4090 | 12GB | 1 x RTX 4090 | 14GB | 1 x RTX 4090 | 12GB | 1 x RTX 4090 | 12GB | 2 x RTX 4090 | 18GB | 1 x RTX 4090 | 18GB |
| 50% | 1 x RTX 4090 | 12GB | 1 x RTX 4090 | 14GB | 2 x RTX 4090 | 12GB | 2 x RTX 4090 | 12GB | 3 x RTX 4090 | 18GB | 2 x RTX 4090 | 18GB |
| 75% | 1 x RTX 4090 | 12GB | 1 x RTX 4090 | 14GB | 2 x RTX 4090 | 12GB | 2 x RTX 4090 | 12GB | 4 x RTX 4090 | 18GB | 2 x RTX 4090 | 18GB |
| All/Last | 1 x RTX 4090 | 12GB | 1 x RTX 4090 | 14GB | 4 x Quadro RTX 6000 | 12GB | 2 x TITAN RTX | 12GB | 4 x Quadro RTX 6000 | 18GB | 2 x TITAN RTX | 18GB |
| ESM2-3B - 1st | 1 x RTX 4090 | 30GB | 1 x RTX 4090 | 14GB | 2 x TITAN RTX | 40GB | 1 x RTX 4090 | 40GB | 2 x TITAN RTX | 18GB | 1 x RTX 4090 | 18GB |
| 25% | 1 x RTX 4090 | 30GB | 1 x RTX 4090 | 14GB | 2 x TITAN RTX | 40GB | 2 x TITAN RTX | 40GB | 2 x TITAN RTX | 18GB | 2 x TITAN RTX | 18GB |
| 50% | 1 x RTX 4090 | 30GB | 1 x RTX 4090 | 14GB | 4 x TITAN RTX | 40GB | 4 x TITAN RTX | 40GB | 4 x TITAN RTX | 18GB | 4 x TITAN RTX | 18GB |
| 75% | 1 x RTX 4090 | 30GB | 1 x RTX 4090 | 14GB | 3 x A100 (40 GiB) | 40GB | 4 x TITAN RTX | 40GB | 3 x A100 (40 GiB) | 18GB | 4 x TITAN RTX | 18GB |
| All/Last | 1 x RTX 4090 | 30GB | 1 x RTX 4090 | 14GB | 4 x A100 (80 GiB) | 40GB | 4 x Quadro RTX 6000 | 40GB | 3 x A100 (40 GiB) | 18GB | 4 x Quadro RTX 6000 | 18GB |
| ProGen2-xlarge - 1st | 1 x RTX 4090 | 60GB | 1 x RTX 4090 | 14GB | 1 x RTX 4090 | 40GB | 1 x RTX 4090 | 40GB | 2 x TITAN RTX | 40GB | 1 x RTX 4090 | 40GB |
| 25% | 1 x RTX 4090 | 60GB | 1 x RTX 4090 | 14GB | 2 x TITAN RTX | 40GB | 2 x TITAN RTX | 40GB | 2 x TITAN RTX | 40GB | 2 x TITAN RTX | 40GB |
| 50% | 1 x RTX 4090 | 60GB | 1 x RTX 4090 | 14GB | 4 x TITAN RTX | 40GB | 4 x TITAN RTX | 40GB | 4 x TITAN RTX | 40GB | 4 x TITAN RTX | 40GB |
| 75% | 1 x RTX 4090 | 60GB | 1 x RTX 4090 | 14GB | 4 x A100 (40 GiB) | 40GB | 4 x TITAN RTX | 40GB | 3 x A100 (40 GiB) | 40GB | 4 x TITAN RTX | 40GB |
| All/Last | 1 x A100 (80 GiB) | 60GB | 1 x RTX 4090 | 14GB | 4 x A100 (80 GiB) | 40GB | 4 x Quadro RTX 6000 | 40GB | 4 x A100 (80 GiB) | 40GB | 4 x Quadro RTX 6000 | 40GB |
| ESM2-15B - 1st | 1 x RTX 4090 | 80GB | 1 x RTX 4090 | 14GB | 2 x TITAN RTX | 70GB | 1 x RTX 4090 | 70GB | 2 x TITAN RTX | 75GB | 1 x RTX 4090 | 75GB |
| 25% | 1 x RTX 4090 | 80GB | 1 x RTX 4090 | 14GB | 2 x TITAN RTX | 70GB | 2 x TITAN RTX | 70GB | 3 x TITAN RTX | 75GB | 4 x TITAN RTX | 75GB |
| 50% | 1 x RTX 4090 | 80GB | 1 x RTX 4090 | 14GB | 4 x A100 (40 GiB) | 70GB | 4 x TITAN RTX | 70GB | 4 x A100 (40 GiB) | 75GB | 4 x TITAN RTX | 75GB |
| 75% | 1 x A100 (80 GiB) | 80GB | 1 x RTX 4090 | 14GB | 4 x A100 (80 GiB) | 70GB | 4 x TITAN RTX | 70GB | 4 x A100 (80 GiB) | 75GB | 4 x TITAN RTX | 75GB |
| All/Last | 1 x A100 (80 GiB) | 80GB | 1 x RTX 4090 | 14GB | 4 x A100 (80 GiB) | 70GB | 4 x TITAN RTX | 70GB | 4 x A100 (80 GiB) | 75GB | 4 x A100 (80 GiB) | 75GB |

**Table S7.Detailed computational resources (number of GPUs and type, CPU RAM) used for each TL-based model on *Meltome-mixed* task.**

| PLM + Layer | Embeddings Extraction | | Feature Extraction | | LoRA | | LoRA- | | Adapters | | Adapters- | |
|---|---|---|---|---|---|---|---|---|---|---|---|---|
| | GPUs | CPU RAM | GPUs | CPU RAM | GPUs | CPU RAM | GPUs | CPU RAM | GPUs | CPU RAM | GPUs | CPU RAM |
| ProteinBERT - 1st | 1 x RTX 4090 | 12GB | 1 x RTX 4090 | 10GB | 1 x RTX 2080 Ti | 12GB | 1 x RTX 2080 Ti | 12GB | 1 x RTX 2080 Ti | 8GB | 1 x RTX 2080 Ti | 8GB |
| 25% | 1 x RTX 4090 | 12GB | 1 x RTX 4090 | 10GB | 1 x RTX 2080 Ti | 12GB | 1 x RTX 2080 Ti | 12GB | 1 x RTX 2080 Ti | 8GB | 1 x RTX 2080 Ti | 8GB |
| 50% | 1 x RTX 4090 | 12GB | 1 x RTX 4090 | 10GB | 1 x RTX 2080 Ti | 12GB | 1 x RTX 2080 Ti | 12GB | 1 x RTX 2080 Ti | 8GB | 1 x RTX 2080 Ti | 8GB |
| 75% | 1 x RTX 4090 | 12GB | 1 x RTX 4090 | 10GB | 1 x RTX 4090 | 12GB | 1 x RTX 2080 Ti | 12GB | 1 x RTX 4090 | 8GB | 1 x RTX 2080 Ti | 8GB |
| All/Last | 1 x RTX 4090 | 12GB | 1 x RTX 4090 | 10GB | 1 x RTX 4090 | 12GB | 1 x RTX 4090 | 12GB | 1 x RTX 4090 | 8GB | 1 x RTX 4090 | 8GB |
| ProGen2-small - 1st | 1 x RTX 4090 | 12GB | 1 x RTX 4090 | 10GB | 1 x RTX 4090 | 12GB | 1 x RTX 4090 | 12GB | 1 x RTX 4090 | 12GB | 1 x RTX 4090 | 12GB |
| 25% | 1 x RTX 4090 | 12GB | 1 x RTX 4090 | 10GB | 1 x RTX 4090 | 12GB | 1 x RTX 4090 | 12GB | 1 x RTX 4090 | 12GB | 1 x RTX 4090 | 12GB |
| 50% | 1 x RTX 4090 | 12GB | 1 x RTX 4090 | 10GB | 1 x RTX 4090 | 12GB | 1 x RTX 4090 | 12GB | 1 x RTX 4090 | 12GB | 1 x RTX 4090 | 12GB |
| 75% | 1 x RTX 4090 | 12GB | 1 x RTX 4090 | 10GB | 1 x RTX 4090 | 12GB | 1 x RTX 4090 | 12GB | 1 x RTX 4090 | 12GB | 1 x RTX 4090 | 12GB |
| All/Last | 1 x RTX 4090 | 12GB | 1 x RTX 4090 | 10GB | 1 x RTX 4090 | 12GB | 1 x RTX 4090 | 12GB | 1 x RTX 4090 | 12GB | 1 x RTX 4090 | 12GB |
| ESM2-650M - 1st | 1 x RTX 4090 | 12GB | 1 x RTX 4090 | 14GB | 1 x RTX 3090 | 12GB | 1 x RTX 3090 | 12GB | 1 x RTX 3090 | 18GB | 1 x RTX 3090 | 18GB |
| 25% | 1 x RTX 4090 | 12GB | 1 x RTX 4090 | 14GB | 1 x RTX 3090 | 12GB | 1 x RTX 3090 | 12GB | 1 x RTX 3090 | 18GB | 1 x RTX 3090 | 18GB |
| 50% | 1 x RTX 4090 | 12GB | 1 x RTX 4090 | 14GB | 2 x RTX 4090 | 12GB | 2 x RTX 4090 | 12GB | 2 x RTX 4090 | 18GB | 2 x RTX 4090 | 18GB |
| 75% | 1 x RTX 4090 | 12GB | 1 x RTX 4090 | 14GB | 2 x Quadro RTX 6000 | 12GB | 2 x Quadro RTX 6000 | 12GB | 2 x Quadro RTX 6000 | 18GB | 2 x Quadro RTX 6000 | 18GB |
| All/Last | 1 x RTX 4090 | 12GB | 1 x RTX 4090 | 14GB | 2 x Quadro RTX 6000 | 12GB | 2 x Quadro RTX 6000 | 12GB | 2 x Quadro RTX 6000 | 18GB | 2 x Quadro RTX 6000 | 18GB |
| ProGen2-medium - 1st | 1 x RTX 4090 | 12GB | 1 x RTX 4090 | 14GB | 1 x RTX 3090 | 12GB | 1 x RTX 3090 | 12GB | 1 x RTX 3090 | 18GB | 1 x RTX 3090 | 18GB |
| 25% | 1 x RTX 4090 | 12GB | 1 x RTX 4090 | 14GB | 1 x RTX 3090 | 12GB | 1 x RTX 3090 | 12GB | 1 x RTX 3090 | 18GB | 1 x RTX 3090 | 18GB |
| 50% | 1 x RTX 4090 | 12GB | 1 x RTX 4090 | 14GB | 2 x RTX 4090 | 12GB | 2 x RTX 4090 | 12GB | 2 x RTX 4090 | 18GB | 2 x RTX 4090 | 18GB |
| 75% | 1 x RTX 4090 | 12GB | 1 x RTX 4090 | 14GB | 2 x Quadro RTX 6000 | 12GB | 2 x RTX 4090 | 12GB | 2 x Quadro RTX 6000 | 18GB | 2 x RTX 4090 | 18GB |
| All/Last | 1 x RTX 4090 | 12GB | 1 x RTX 4090 | 14GB | 2 x Quadro RTX 6000 | 12GB | 2 x Quadro RTX 6000 | 12GB | 2 x Quadro RTX 6000 | 18GB | 2 x Quadro RTX 6000 | 18GB |
| ESM2-3B - 1st | 1 x RTX 4090 | 30GB | 1 x RTX 4090 | 14GB | 1 x RTX 3090 | 40GB | 1 x RTX 3090 | 40GB | 1 x RTX 3090 | 18GB | 1 x RTX 3090 | 18GB |
| 25% | 1 x RTX 4090 | 30GB | 1 x RTX 4090 | 14GB | 2 x RTX 4090 | 40GB | 2 x RTX 4090 | 40GB | 2 x RTX 4090 | 18GB | 2 x RTX 4090 | 18GB |
| 50% | 1 x RTX 4090 | 30GB | 1 x RTX 4090 | 14GB | 2 x Quadro RTX 6000 | 40GB | 2 x Quadro RTX 6000 | 40GB | 2 x Quadro RTX 6000 | 18GB | 2 x Quadro RTX 6000 | 18GB |
| 75% | 1 x RTX 4090 | 30GB | 1 x RTX 4090 | 14GB | 3 x Quadro RTX 6000 | 40GB | 3 x Quadro RTX 6000 | 40GB | 3 x Quadro RTX 6000 | 18GB | 2 x Quadro RTX 6000 | 18GB |
| All/Last | 1 x RTX 4090 | 30GB | 1 x RTX 4090 | 14GB | 3 x A100 (40 GiB) | 40GB | 3 x Quadro RTX 6000 | 40GB | 3 x A100 (40 GiB) | 18GB | 2 x Quadro RTX 6000 | 18GB |
| ProGen2-xlarge - 1st | 1 x RTX 4090 | 60GB | 1 x RTX 4090 | 14GB | 1 x RTX 2080 Ti | 40GB | 1 x RTX 2080 Ti | 40GB | 1 x RTX 2080 Ti | 40GB | 1 x RTX 2080 Ti | 40GB |
| 25% | 1 x RTX 4090 | 60GB | 1 x RTX 4090 | 14GB | 2 x RTX 3090 | 40GB | 2 x RTX 3090 | 40GB | 2 x RTX 3090 | 40GB | 2 x RTX 3090 | 40GB |
| 50% | 1 x RTX 4090 | 60GB | 1 x RTX 4090 | 14GB | 2 x RTX 3090 | 40GB | 2 x RTX 3090 | 40GB | 2 x RTX 3090 | 40GB | 2 x RTX 3090 | 40GB |
| 75% | 1 x RTX 4090 | 60GB | 1 x RTX 4090 | 14GB | 2 x A100 (40 GiB) | 40GB | 2 x RTX 3090 | 40GB | 2 x Quadro RTX 6000 | 40GB | 2 x RTX 3090 | 40GB |
| All/Last | 1 x A100 (80 GiB) | 60GB | 1 x RTX 4090 | 14GB | 3 x A100 (80 GiB) | 40GB | 3 x Quadro RTX 6000 | 40GB | 3 x Quadro RTX 6000 | 40GB | 2 x Quadro RTX 6000 | 40GB |
| ESM2-15B - 1st | 1 x RTX 4090 | 80GB | 1 x RTX 4090 | 14GB | 1 x RTX 2080 Ti | 70GB | 1 x RTX 2080 Ti | 70GB | 1 x RTX 2080 Ti | 75GB | 1 x RTX 2080 Ti | 75GB |
| 25% | 1 x RTX 4090 | 80GB | 1 x RTX 4090 | 14GB | 2 x RTX 3090 | 70GB | 2 x RTX 3090 | 70GB | 3 x RTX 3090 | 75GB | 1 x RTX 3090 | 75GB |
| 50% | 1 x RTX 4090 | 80GB | 1 x RTX 4090 | 14GB | 2 x A100 (40 GiB) | 70GB | 2 x Quadro RTX 6000 | 70GB | 3 x Quadro RTX 6000 | 75GB | 2 x Quadro RTX 6000 | 75GB |
| 75% | 1 x A100 (80 GiB) | 80GB | 1 x RTX 4090 | 14GB | 3 x A100 (80 GiB) | 70GB | 2 x Quadro RTX 6000 | 70GB | 3 x A100 (80 GiB) | 75GB | 2 x Quadro RTX 6000 | 75GB |
| All/Last | 1 x A100 (80 GiB) | 80GB | 1 x RTX 4090 | 14GB | 4 x A100 (80 GiB) | 70GB | 2 x A100 (40 GiB) | 70GB | 3 x A100 (80 GiB) | 75GB | 2 x A100 (80 GiB) | 75GB |

**Table S8. Detailed computational resources (number of GPUs and type, CPU RAM) used for each TL-based model on *RBD-one vs. rest* task.**

| PLM + Layer | Embeddings Extraction | | Feature Extraction | | LoRA | | LoRA- | | Adapters | | Adapters- | |
|---|---|---|---|---|---|---|---|---|---|---|---|---|
| | GPUs | CPU RAM | GPUs | CPU RAM | GPUs | CPU RAM | GPUs | CPU RAM | GPUs | CPU RAM | GPUs | CPU RAM |
| ProteinBERT - 1st | 1 x RTX 4090 | 12GB | 1 x RTX 4090 | 10GB | 1 x RTX 2080 Ti | 12GB | 1 x RTX 2080 Ti | 12GB | 1 x RTX 2080 Ti | 8GB | 1 x RTX 2080 Ti | 8GB |
| 25% | 1 x RTX 4090 | 12GB | 1 x RTX 4090 | 10GB | 1 x RTX 2080 Ti | 12GB | 1 x RTX 2080 Ti | 12GB | 1 x RTX 2080 Ti | 8GB | 1 x RTX 2080 Ti | 8GB |
| 50% | 1 x RTX 4090 | 12GB | 1 x RTX 4090 | 10GB | 1 x RTX 2080 Ti | 12GB | 1 x RTX 2080 Ti | 12GB | 1 x RTX 2080 Ti | 8GB | 1 x RTX 2080 Ti | 8GB |
| 75% | 1 x RTX 4090 | 12GB | 1 x RTX 4090 | 10GB | 1 x RTX 2080 Ti | 12GB | 1 x RTX 2080 Ti | 12GB | 1 x RTX 2080 Ti | 8GB | 1 x RTX 2080 Ti | 8GB |
| All/Last | 1 x RTX 4090 | 12GB | 1 x RTX 4090 | 10GB | 1 x RTX 4090 | 12GB | 1 x RTX 4090 | 12GB | 1 x RTX 4090 | 8GB | 1 x RTX 4090 | 8GB |
| ProGen2-small - 1st | 1 x RTX 4090 | 12GB | 1 x RTX 4090 | 10GB | 1 x RTX 4090 | 12GB | 1 x RTX 4090 | 12GB | 1 x RTX 4090 | 12GB | 1 x RTX 4090 | 12GB |
| 25% | 1 x RTX 4090 | 12GB | 1 x RTX 4090 | 10GB | 1 x RTX 4090 | 12GB | 1 x RTX 4090 | 12GB | 1 x RTX 4090 | 12GB | 1 x RTX 4090 | 12GB |
| 50% | 1 x RTX 4090 | 12GB | 1 x RTX 4090 | 10GB | 1 x RTX 4090 | 12GB | 1 x RTX 4090 | 12GB | 1 x RTX 4090 | 12GB | 1 x RTX 4090 | 12GB |
| 75% | 1 x RTX 4090 | 12GB | 1 x RTX 4090 | 10GB | 1 x RTX 4090 | 12GB | 1 x RTX 4090 | 12GB | 1 x RTX 4090 | 12GB | 1 x RTX 4090 | 12GB |
| All/Last | 1 x RTX 4090 | 12GB | 1 x RTX 4090 | 10GB | 1 x RTX 4090 | 12GB | 1 x RTX 4090 | 12GB | 1 x RTX 4090 | 12GB | 1 x RTX 4090 | 12GB |
| ESM2-650M - 1st | 1 x RTX 4090 | 12GB | 1 x RTX 4090 | 14GB | 1 x RTX 3090 | 12GB | 1 x RTX 3090 | 12GB | 1 x RTX 3090 | 18GB | 1 x RTX 3090 | 18GB |
| 25% | 1 x RTX 4090 | 12GB | 1 x RTX 4090 | 14GB | 1 x RTX 3090 | 12GB | 1 x RTX 3090 | 12GB | 1 x RTX 3090 | 18GB | 1 x RTX 3090 | 18GB |
| 50% | 1 x RTX 4090 | 12GB | 1 x RTX 4090 | 14GB | 2 x RTX 4090 | 12GB | 2 x RTX 4090 | 12GB | 2 x RTX 4090 | 18GB | 2 x RTX 4090 | 18GB |
| 75% | 1 x RTX 4090 | 12GB | 1 x RTX 4090 | 14GB | 2 x Quadro RTX 6000 | 12GB | 2 x Quadro RTX 6000 | 12GB | 2 x Quadro RTX 6000 | 18GB | 2 x Quadro RTX 6000 | 18GB |
| All/Last | 1 x RTX 4090 | 12GB | 1 x RTX 4090 | 14GB | 2 x Quadro RTX 6000 | 12GB | 2 x Quadro RTX 6000 | 12GB | 2 x Quadro RTX 6000 | 18GB | 2 x Quadro RTX 6000 | 18GB |
| ProGen2-medium - 1st | 1 x RTX 4090 | 12GB | 1 x RTX 4090 | 14GB | 1 x RTX 3090 | 12GB | 1 x RTX 4090 | 12GB | 1 x RTX 3090 | 18GB | 1 x RTX 4090 | 18GB |
| 25% | 1 x RTX 4090 | 12GB | 1 x RTX 4090 | 14GB | 1 x RTX 3090 | 12GB | 1 x RTX 3090 | 12GB | 1 x RTX 3090 | 18GB | 1 x RTX 3090 | 18GB |
| 50% | 1 x RTX 4090 | 12GB | 1 x RTX 4090 | 14GB | 2 x RTX 4090 | 12GB | 2 x RTX 4090 | 12GB | 2 x RTX 4090 | 18GB | 2 x RTX 4090 | 18GB |
| 75% | 1 x RTX 4090 | 12GB | 1 x RTX 4090 | 14GB | 2 x Quadro RTX 6000 | 12GB | 2 x Quadro RTX 6000 | 12GB | 2 x Quadro RTX 6000 | 18GB | 2 x Quadro RTX 6000 | 18GB |
| All/Last | 1 x RTX 4090 | 12GB | 1 x RTX 4090 | 14GB | 2 x Quadro RTX 6000 | 12GB | 2 x Quadro RTX 6000 | 12GB | 2 x Quadro RTX 6000 | 18GB | 2 x Quadro RTX 6000 | 18GB |
| ESM2-3B - 1st | 1 x RTX 4090 | 30GB | 1 x RTX 4090 | 14GB | 1 x RTX 3090 | 40GB | 1 x RTX 3090 | 40GB | 1 x RTX 3090 | 18GB | 1 x RTX 3090 | 18GB |
| 25% | 1 x RTX 4090 | 30GB | 1 x RTX 4090 | 14GB | 1 x RTX 4090 | 40GB | 1 x RTX 4090 | 40GB | 1 x RTX 4090 | 18GB | 1 x RTX 4090 | 18GB |
| 50% | 1 x RTX 4090 | 30GB | 1 x RTX 4090 | 14GB | 1 x Quadro RTX 6000 | 40GB | 1 x Quadro RTX 6000 | 40GB | 1 x Quadro RTX 6000 | 18GB | 1 x Quadro RTX 6000 | 18GB |
| 75% | 1 x RTX 4090 | 30GB | 1 x RTX 4090 | 14GB | 1 x Quadro RTX 6000 | 40GB | 1 x Quadro RTX 6000 | 40GB | 1 x Quadro RTX 6000 | 18GB | 1 x Quadro RTX 6000 | 18GB |
| All/Last | 1 x RTX 4090 | 30GB | 1 x RTX 4090 | 14GB | 1 x A100 (40 GiB) | 40GB | 1 x Quadro RTX 6000 | 40GB | 1 x A100 (40 GiB) | 18GB | 1 x Quadro RTX 6000 | 18GB |
| ProGen2-xlarge - 1st | 1 x RTX 4090 | 60GB | 1 x RTX 4090 | 14GB | 1 x RTX 2080 Ti | 40GB | 1 x RTX 2080 Ti | 40GB | 1 x RTX 2080 Ti | 40GB | 1 x RTX 2080 Ti | 40GB |
| 25% | 1 x RTX 4090 | 60GB | 1 x RTX 4090 | 14GB | 1 x RTX 3090 | 40GB | 1 x RTX 3090 | 40GB | 1 x RTX 3090 | 40GB | 1 x RTX 3090 | 40GB |
| 50% | 1 x RTX 4090 | 60GB | 1 x RTX 4090 | 14GB | 2 x RTX 3090 | 40GB | 2 x RTX 3090 | 40GB | 2 x RTX 3090 | 40GB | 1 x RTX 3090 | 40GB |
| 75% | 1 x RTX 4090 | 60GB | 1 x RTX 4090 | 14GB | 2 x A100 (40 GiB) | 40GB | 2 x RTX 3090 | 40GB | 2 x A100 (40 GiB) | 40GB | 1 x RTX 3090 | 40GB |
| All/Last | 1 x A100 (80 GiB) | 60GB | 1 x RTX 4090 | 14GB | 2 x A100 (80 GiB) | 40GB | 2 x Quadro RTX 6000 | 40GB | 2 x A100 (80 GiB) | 40GB | 1 x Quadro RTX 6000 | 40GB |
| ESM2-15B - 1st | 1 x RTX 4090 | 80GB | 1 x RTX 4090 | 14GB | 1 x RTX 2080 Ti | 70GB | 1 x RTX 2080 Ti | 70GB | 1 x RTX 2080 Ti | 75GB | 1 x RTX 2080 Ti | 75GB |
| 25% | 1 x RTX 4090 | 80GB | 1 x RTX 4090 | 14GB | 2 x RTX 3090 | 70GB | 2 x RTX 3090 | 70GB | 2 x RTX 3090 | 75GB | 2 x RTX 3090 | 75GB |
| 50% | 1 x RTX 4090 | 80GB | 1 x RTX 4090 | 14GB | 2 x A100 (40 GiB) | 70GB | 2 x RTX 3090 | 70GB | 2 x A100 (40 GiB) | 75GB | 2 x RTX 3090 | 75GB |
| 75% | 1 x A100 (80 GiB) | 80GB | 1 x RTX 4090 | 14GB | 3 x A100 (80 GiB) | 70GB | 2 x Quadro RTX 6000 | 70GB | 3 x A100 (80 GiB) | 75GB | 2 x RTX 4090 | 75GB |
| All/Last | 1 x A100 (80 GiB) | 80GB | 1 x RTX 4090 | 14GB | 3 x A100 (80 GiB) | 70GB | 3 x Quadro RTX 6000 | 70GB | 3 x A100 (80 GiB) | 75GB | 2 x A100 (80 GiB) | 75GB |

**Table S9. Detailed computational resources (number of GPUs and type, CPU RAM) used for each TL-based model on *Trastuzumab-one vs. rest* task.**

| PLM + Layer | Embeddings Extraction | | Feature Extraction | | LoRA | | LoRA- | | Adapters | | Adapters- | |
|---|---|---|---|---|---|---|---|---|---|---|---|---|
| | GPUs | CPU RAM | GPUs | CPU RAM | GPUs | CPU RAM | GPUs | CPU RAM | GPUs | CPU RAM | GPUs | CPU RAM |
| ProteinBERT - 1st | 1 x RTX 4090 | 35GB | 1 x RTX 4090 | 35GB | 1 x RTX 2080 Ti | 12GB | 1 x RTX 2080 Ti | 12GB | 1 x RTX 2080 Ti | 8GB | 1 x RTX 2080 Ti | 8GB |
| 25% | 1 x RTX 4090 | 35GB | 1 x RTX 4090 | 35GB | 1 x RTX 2080 Ti | 12GB | 1 x RTX 2080 Ti | 12GB | 1 x RTX 2080 Ti | 8GB | 1 x RTX 2080 Ti | 8GB |
| 50% | 1 x RTX 4090 | 35GB | 1 x RTX 4090 | 35GB | 1 x RTX 2080 Ti | 12GB | 1 x RTX 2080 Ti | 12GB | 1 x RTX 2080 Ti | 8GB | 1 x RTX 2080 Ti | 8GB |
| 75% | 1 x RTX 4090 | 35GB | 1 x RTX 4090 | 35GB | 1 x RTX 4090 | 12GB | 1 x RTX 2080 Ti | 12GB | 1 x RTX 4090 | 8GB | 1 x RTX 2080 Ti | 8GB |
| All/Last | 1 x RTX 4090 | 35GB | 1 x RTX 4090 | 35GB | 1 x RTX 4090 | 12GB | 1 x RTX 4090 | 12GB | 1 x RTX 4090 | 8GB | 1 x RTX 4090 | 8GB |
| ProGen2-small - 1st | 1 x RTX 4090 | 55GB | 1 x RTX 4090 | 55GB | 1 x RTX 4090 | 12GB | 1 x RTX 4090 | 12GB | 1 x RTX 4090 | 12GB | 1 x RTX 4090 | 12GB |
| 25% | 1 x RTX 4090 | 55GB | 1 x RTX 4090 | 55GB | 1 x RTX 4090 | 12GB | 1 x RTX 4090 | 12GB | 1 x RTX 4090 | 12GB | 1 x RTX 4090 | 12GB |
| 50% | 1 x RTX 4090 | 55GB | 1 x RTX 4090 | 55GB | 1 x RTX 4090 | 12GB | 1 x RTX 4090 | 12GB | 1 x RTX 4090 | 12GB | 1 x RTX 4090 | 12GB |
| 75% | 1 x RTX 4090 | 55GB | 1 x RTX 4090 | 55GB | 1 x RTX 4090 | 12GB | 1 x RTX 4090 | 12GB | 1 x RTX 4090 | 12GB | 1 x RTX 4090 | 12GB |
| All/Last | 1 x RTX 4090 | 55GB | 1 x RTX 4090 | 55GB | 1 x RTX 4090 | 12GB | 1 x RTX 4090 | 12GB | 1 x RTX 4090 | 12GB | 1 x RTX 4090 | 12GB |
| ESM2-650M - 1st | 1 x RTX 4090 | 60GB | 1 x RTX 4090 | 60GB | 1 x RTX 3090 | 12GB | 1 x RTX 3090 | 12GB | 1 x RTX 3090 | 18GB | 1 x RTX 3090 | 18GB |
| 25% | 1 x RTX 4090 | 60GB | 1 x RTX 4090 | 60GB | 1 x RTX 3090 | 12GB | 1 x RTX 3090 | 12GB | 1 x RTX 3090 | 18GB | 1 x RTX 3090 | 18GB |
| 50% | 1 x RTX 4090 | 60GB | 1 x RTX 4090 | 60GB | 2 x RTX 4090 | 12GB | 2 x RTX 4090 | 12GB | 2 x RTX 4090 | 18GB | 2 x RTX 4090 | 18GB |
| 75% | 1 x RTX 4090 | 60GB | 1 x RTX 4090 | 60GB | 2 x Quadro RTX 6000 | 12GB | 2 x Quadro RTX 6000 | 12GB | 2 x Quadro RTX 6000 | 18GB | 2 x Quadro RTX 6000 | 18GB |
| All/Last | 1 x RTX 4090 | 60GB | 1 x RTX 4090 | 60GB | 2 x Quadro RTX 6000 | 12GB | 2 x Quadro RTX 6000 | 12GB | 2 x Quadro RTX 6000 | 18GB | 2 x Quadro RTX 6000 | 18GB |
| ProGen2-medium - 1st | 1 x RTX 4090 | 70GB | 1 x RTX 4090 | 70GB | 1 x RTX 3090 | 12GB | 1 x RTX 3090 | 12GB | 1 x RTX 3090 | 18GB | 1 x RTX 3090 | 18GB |
| 25% | 1 x RTX 4090 | 70GB | 1 x RTX 4090 | 70GB | 1 x RTX 3090 | 12GB | 1 x RTX 3090 | 12GB | 1 x RTX 3090 | 18GB | 1 x RTX 3090 | 18GB |
| 50% | 1 x RTX 4090 | 70GB | 1 x RTX 4090 | 70GB | 2 x RTX 4090 | 12GB | 2 x RTX 4090 | 12GB | 2 x RTX 4090 | 18GB | 2 x RTX 4090 | 18GB |
| 75% | 1 x RTX 4090 | 70GB | 1 x RTX 4090 | 70GB | 2 x Quadro RTX 6000 | 12GB | 2 x RTX 4090 | 12GB | 2 x Quadro RTX 6000 | 18GB | 2 x RTX 4090 | 18GB |
| All/Last | 1 x RTX 4090 | 70GB | 1 x RTX 4090 | 70GB | 2 x Quadro RTX 6000 | 12GB | 2 x Quadro RTX 6000 | 12GB | 2 x Quadro RTX 6000 | 18GB | 2 x Quadro RTX 6000 | 18GB |
| ESM2-3B - 1st | 1 x RTX 4090 | 120GB | 1 x RTX 4090 | 120GB | 1 x RTX 3090 | 40GB | 1 x RTX 3090 | 40GB | 1 x RTX 3090 | 18GB | 1 x RTX 3090 | 18GB |
| 25% | 1 x RTX 4090 | 120GB | 1 x RTX 4090 | 120GB | 2 x RTX 4090 | 40GB | 2 x RTX 4090 | 40GB | 2 x RTX 4090 | 18GB | 2 x RTX 4090 | 18GB |
| 50% | 1 x RTX 4090 | 120GB | 1 x RTX 4090 | 120GB | 2 x Quadro RTX 6000 | 40GB | 2 x Quadro RTX 6000 | 40GB | 2 x Quadro RTX 6000 | 18GB | 2 x Quadro RTX 6000 | 18GB |
| 75% | 1 x RTX 4090 | 120GB | 1 x RTX 4090 | 120GB | 3 x Quadro RTX 6000 | 40GB | 3 x Quadro RTX 6000 | 40GB | 3 x Quadro RTX 6000 | 18GB | 3 x Quadro RTX 6000 | 18GB |
| All/Last | 1 x RTX 4090 | 120GB | 1 x RTX 4090 | 120GB | 3 x A100 (40 GiB) | 40GB | 3 x Quadro RTX 6000 | 40GB | 3 x A100 (40 GiB) | 18GB | 3 x Quadro RTX 6000 | 18GB |
| ProGen2-xlarge - 1st | 1 x RTX 4090 | 200GB | 1 x RTX 4090 | 200GB | 1 x RTX 2080 Ti | 40GB | 1 x RTX 2080 Ti | 40GB | 1 x RTX 2080 Ti | 40GB | 1 x RTX 2080 Ti | 40GB |
| 25% | 1 x RTX 4090 | 200GB | 1 x RTX 4090 | 200GB | 2 x RTX 3090 | 40GB | 2 x RTX 3090 | 40GB | 2 x RTX 3090 | 40GB | 2 x RTX 3090 | 40GB |
| 50% | 1 x RTX 4090 | 200GB | 1 x RTX 4090 | 200GB | 2 x RTX 3090 | 40GB | 2 x RTX 3090 | 40GB | 2 x RTX 3090 | 40GB | 2 x RTX 3090 | 40GB |
| 75% | 1 x RTX 4090 | 200GB | 1 x RTX 4090 | 200GB | 2 x A100 (40 GiB) | 40GB | 2 x RTX 3090 | 40GB | 2 x A100 (40 GiB) | 40GB | 2 x RTX 3090 | 40GB |
| All/Last | 1 x RTX 4090 | 200GB | 1 x RTX 4090 | 200GB | 3 x A100 (80 GiB) | 40GB | 3 x Quadro RTX 6000 | 40GB | 3 x A100 (80 GiB) | 40GB | 3 x Quadro RTX 6000 | 40GB |
| ESM2-15B - 1st | 1 x RTX 4090 | 250GB | 1 x RTX 4090 | 250GB | 1 x RTX 2080 Ti | 70GB | 1 x RTX 2080 Ti | 70GB | 1 x RTX 2080 Ti | 75GB | 1 x RTX 2080 Ti | 75GB |
| 25% | 1 x RTX 4090 | 250GB | 1 x RTX 4090 | 250GB | 4 x RTX 3090 | 70GB | 4 x RTX 3090 | 70GB | 4 x RTX 3090 | 75GB | 4 x RTX 3090 | 75GB |
| 50% | 1 x RTX 4090 | 250GB | 1 x RTX 4090 | 250GB | 4 x A100 (40 GiB) | 70GB | 4 x Quadro RTX 6000 | 70GB | 4 x A100 (40 GiB) | 75GB | 4 x Quadro RTX 6000 | 75GB |
| 75% | 1 x RTX 4090 | 250GB | 1 x RTX 4090 | 250GB | 4 x A100 (80 GiB) | 70GB | 4 x A100 (40 GiB) | 70GB | 4 x A100 (80 GiB) | 75GB | 4 x Quadro RTX 6000 | 75GB |
| All/Last | 1 x A100 (80 GiB) | 250GB | 1 x RTX 4090 | 250GB | 4 x A100 (80 GiB) | 70GB | 4 x A100 (80 GiB) | 70GB | 4 x A100 (80 GiB) | 75GB | 4 x A100 (80 GiB) | 75GB |

**Table S10. Detailed computational resources (number of GPUs and type, CPU RAM) used for each TL-based model on *SS3-sampled* task.**

### D. Extended results tables

We present an extended representation of the results, including Table S11 which provides statistical summaries of the box plots in Figure 4B and heatmaps in Figures S1-S8, displaying the result metrics for all 3,150 TL setups evaluated.

| Task | Stat | FE | FT |
|---|---|---|---|
| AAV - sampled | Q1 | -5.51% | 3.30% |
| | Q3 | -2.21% | 5.85% |
| | Median | -3.20% | 4.25% |
| | Max | 1.28% | 7.87% |
| AAV - one vs rest | Q1 | -37.98% | 28.60% |
| | Q3 | -13.79% | 43.79% |
| | Median | -32.34% | 38.03% |
| | Max | 8.45% | 47.17% |
| GB1 - three vs rest | Q1 | -9.42% | -1.47% |
| | Q3 | -5.13% | 4.24% |
| | Median | -5.65% | 2.89% |
| | Max | -4.00% | 4.90% |
| GB1 - one vs rest | Q1 | -0.76% | -8.52% |
| | Q3 | 34.02% | 15.08% |
| | Median | 26.43% | 0.74% |
| | Max | 37.67% | 30.98% |
| Meltome - mixed | Q1 | 37.94% | 61.55% |
| | Q3 | 58.12% | 88.95% |
| | Median | 48.99% | 70.79% |
| | Max | 97.13% | 117.83% |
| SS3 - sampled | Q1 | -10.84% | -5.20% |
| | Q3 | 0.86% | 20.64% |
| | Median | -6.82% | 2.36% |
| | Max | 19.48% | 25.24% |

**Table S11. Boxplot numerical values of Figure 4B in detail**

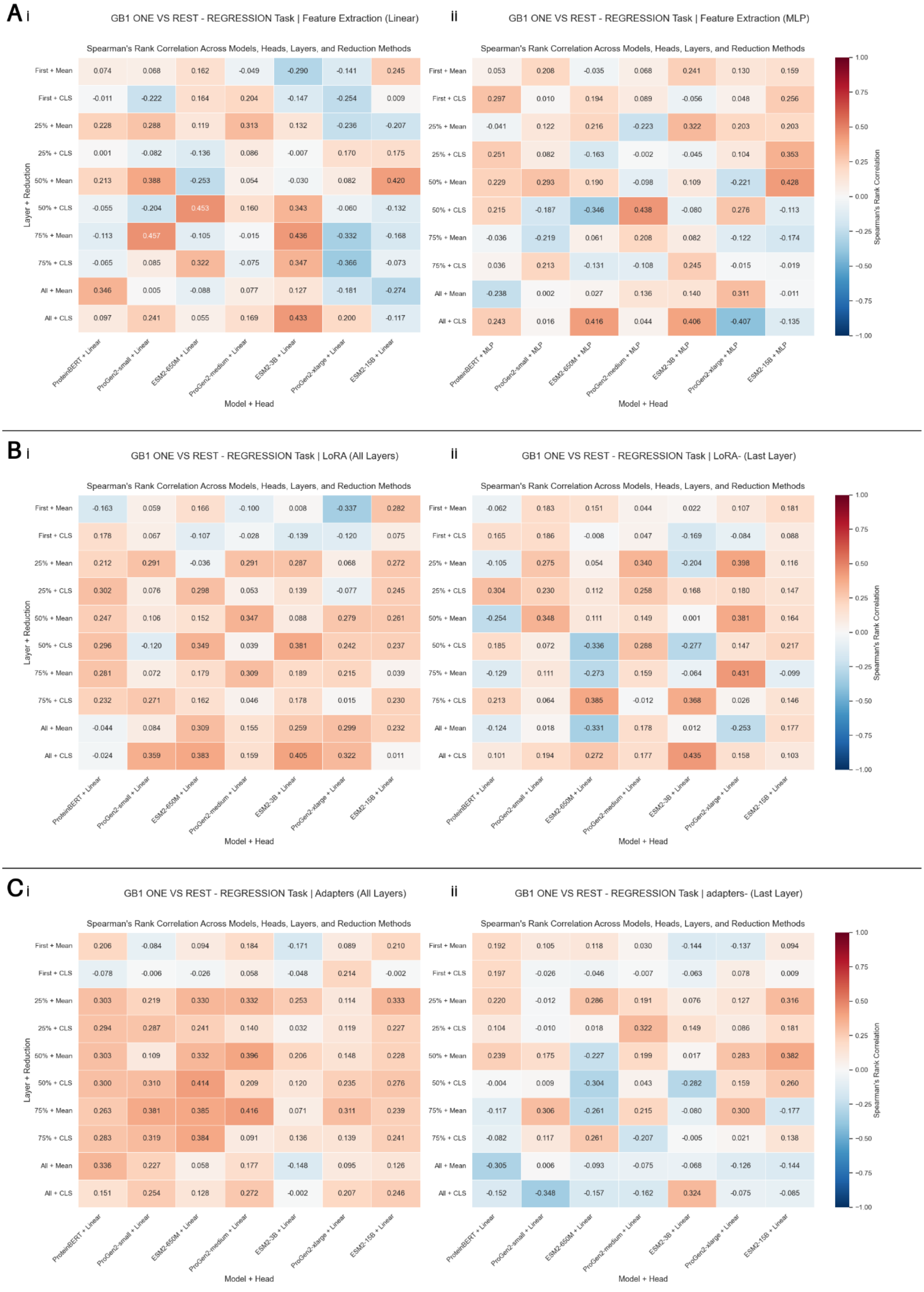


**Figure S1. Detailed results for *GB1-one vs. rest* task.**

Spearman's rank correlation is used as a performance metric for each setup, with x-axis showing the PLM and the head used and the y-axis representing the layers used and the pooling method employed; mean stands for mean pooling and CLS for pooling the classification token for BERT-based PLMs (ESM2, ProteinBERT) and the EOS token for GPT-based PLMs (ProGen2). (A) Feature extraction detailed results using (i) a linear downstream head and (ii) a MLP with one hidden layer as a downstream head. (B) LoRA detailed results when (i) applying LoRA to all layers of PLMs and (ii) applying LoRA to the last layer of PLMs. (C) Adapters

detailed results when (i) applying adapters to all layers of PLMs and (ii) applying adapters to the last layer of PLMs. Empty cells represent work in progress, due to the computational burden these setups bear.

TL: Transfer Learning; PLM: Protein Language Model; LoRA: Low Rank Adaptation

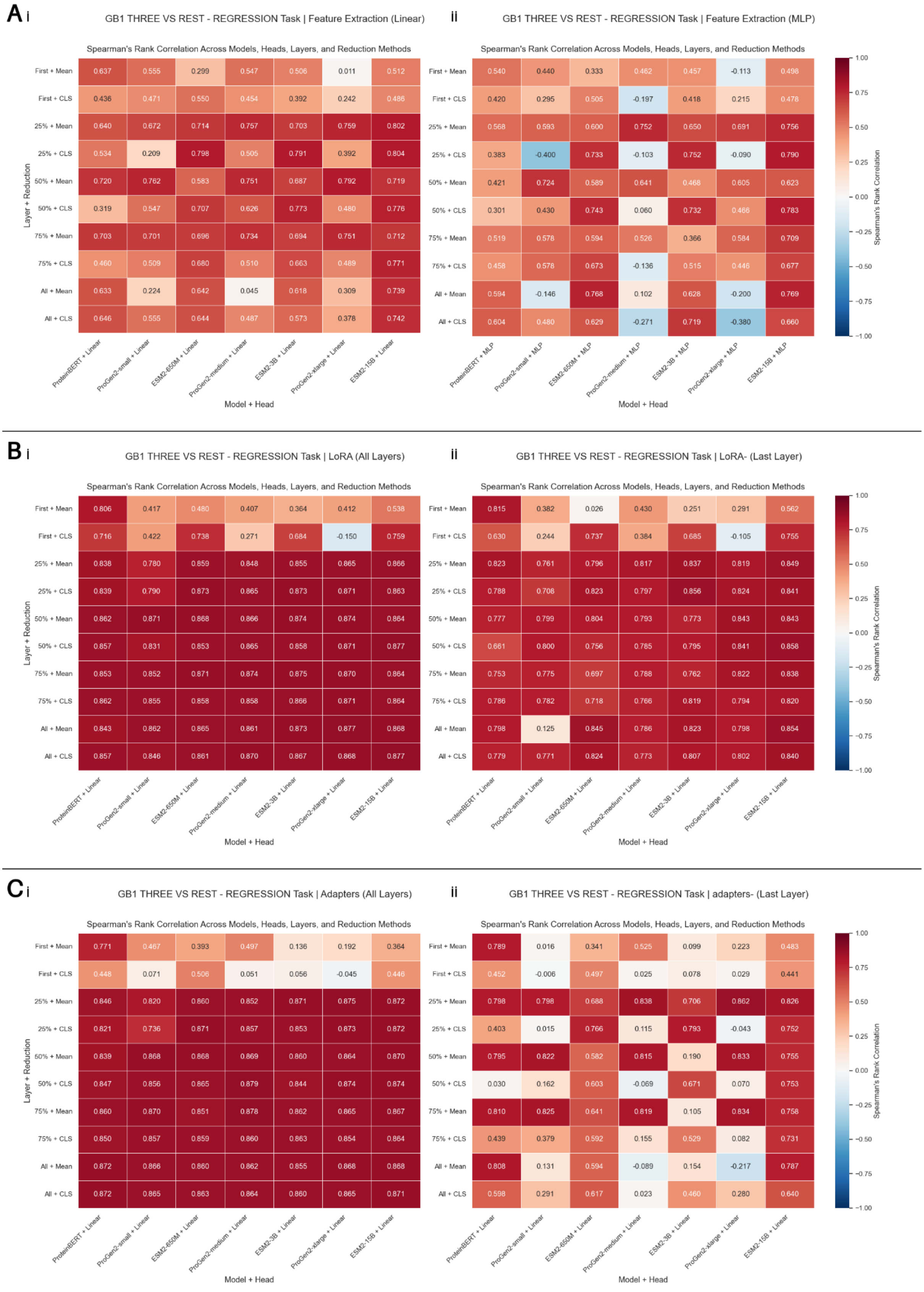


**Figure S2. Detailed results for *GB1-three vs. rest* task.**

Spearman's rank correlation is used as a performance metric for each setup, with x-axis showing the PLM and the head used and the y-axis representing the layers used and the pooling method employed; mean stands for mean pooling and CLS for pooling the classification token for BERT-based PLMs (ESM2, ProteinBERT) and the EOS token for GPT-based PLMs (ProGen2). (A) Feature extraction detailed results using (i) a linear downstream head and (ii) a MLP with one hidden layer as a downstream head. (B) LoRA detailed results when (i) applying LoRA to all layers of PLMs and (ii) applying LoRA to the last layer of PLMs. (C) Adapters

detailed results when (i) applying adapters to all layers of PLMs and (ii) applying adapters to the last layer of PLMs. Empty cells represent work in progress, due to the computational burden these setups bear.

TL: Transfer Learning; PLM: Protein Language Model; LoRA: Low Rank Adaptation

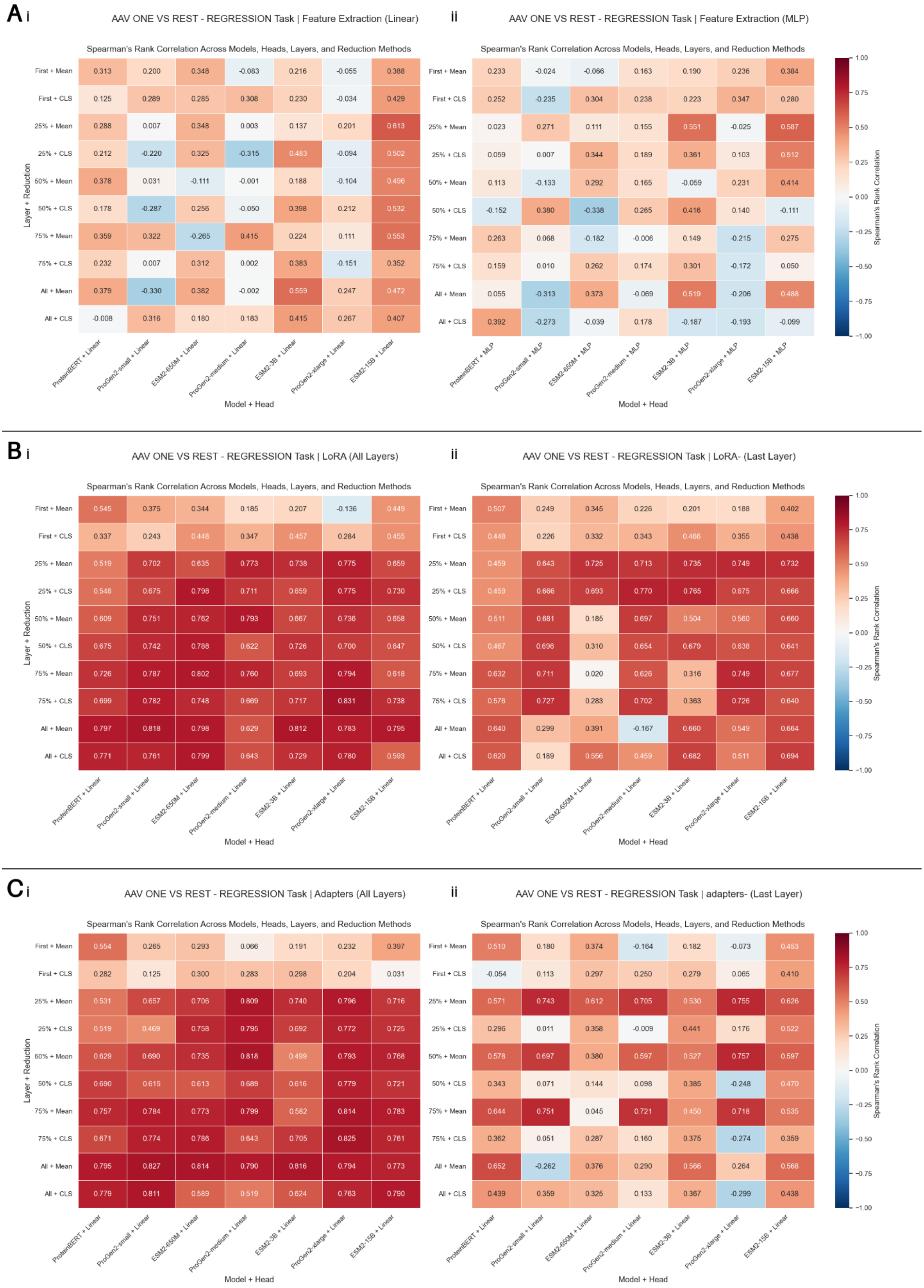


**Figure S3. Detailed results for *AAV-one vs. rest* task.**

Spearman's rank correlation is used as a performance metric for each setup, with x-axis showing the PLM and the head used and the y-axis representing the layers used and the pooling method employed; mean stands for mean pooling and CLS for pooling the classification token for BERT-based PLMs (ESM2, ProteinBERT) and the EOS token for GPT-based PLMs (ProGen2). (A) Feature extraction detailed results using (i) a linear downstream head and (ii) a MLP with one hidden layer as a downstream head. (B) LoRA detailed results when (i) applying LoRA to all layers of PLMs and (ii) applying LoRA to the last layer of PLMs. (C) Adapters

detailed results when (i) applying adapters to all layers of PLMs and (ii) applying adapters to the last layer of PLMs. Empty cells represent work in progress, due to the computational burden these setups bear.

TL: Transfer Learning; PLM: Protein Language Model; LoRA: Low Rank Adaptation

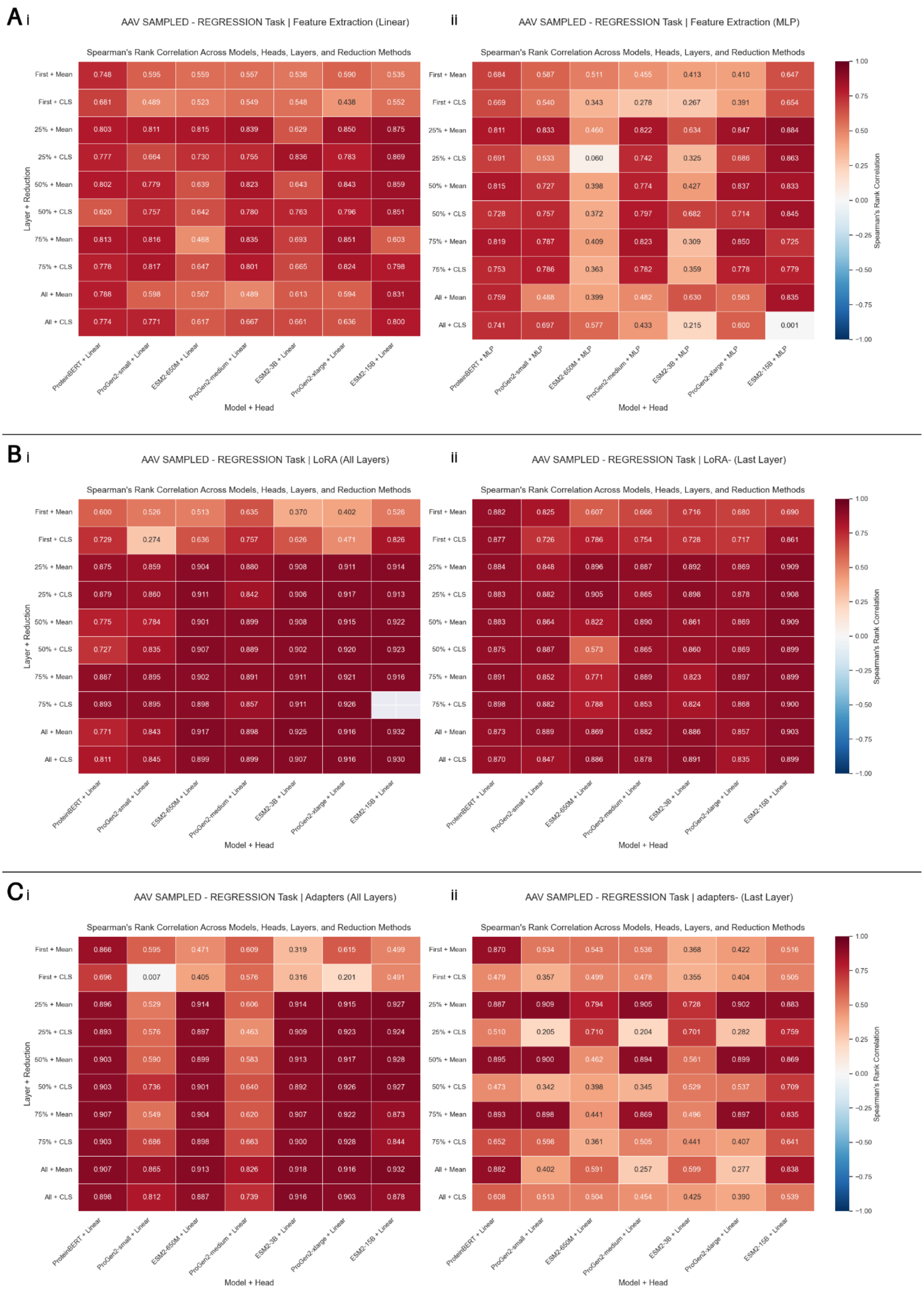


**Figure S4. Detailed results for *AAV-sampled* task.**

Spearman's rank correlation is used as a performance metric for each setup, with x-axis showing the PLM and the head used and the y-axis representing the layers used and the pooling method employed; mean stands for mean pooling and CLS for pooling the classification token for BERT-based PLMs (ESM2, ProteinBERT) and the EOS token for GPT-based PLMs (ProGen2). (A) Feature extraction detailed results using (i) a linear downstream head and (ii) a MLP with one hidden layer as a downstream head. (B) LoRA detailed results when (i) applying LoRA to all layers of PLMs and (ii) applying LoRA to the last layer of PLMs. (C) Adapters

detailed results when (i) applying adapters to all layers of PLMs and (ii) applying adapters to the last layer of PLMs. Empty cells represent work in progress, due to the computational burden these setups bear.

TL: Transfer Learning; PLM: Protein Language Model; LoRA: Low Rank Adaptation

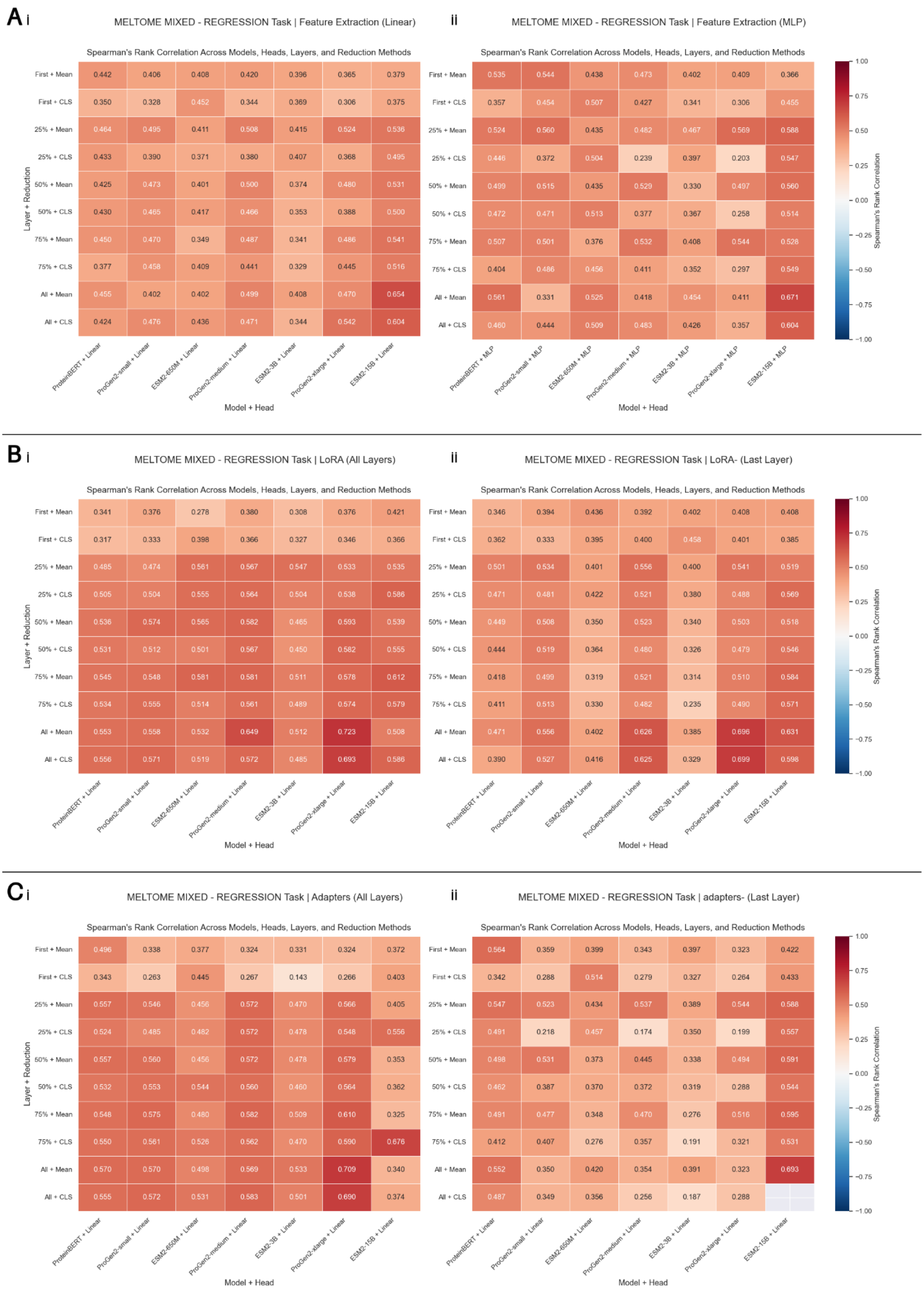


**Figure S5. Detailed results for *Meltome-mixed* task.**

Spearman's rank correlation is used as a performance metric for each setup, with x-axis showing the PLM and the head used and the y-axis representing the layers used and the pooling method employed; mean stands for mean pooling and CLS for pooling the classification token for BERT-based PLMs (ESM2, ProteinBERT) and the EOS token for GPT-based PLMs (ProGen2). (A) Feature extraction detailed results using (i) a linear downstream head and (ii) a MLP with one hidden layer as a downstream head. (B) LoRA detailed results when (i) applying LoRA to all layers of PLMs and (ii) applying LoRA to the last layer of PLMs. (C) Adapters

detailed results when (i) applying adapters to all layers of PLMs and (ii) applying adapters to the last layer of PLMs. Empty cells represent work in progress, due to the computational burden these setups bear.

TL: Transfer Learning; PLM: Protein Language Model; LoRA: Low Rank Adaptation

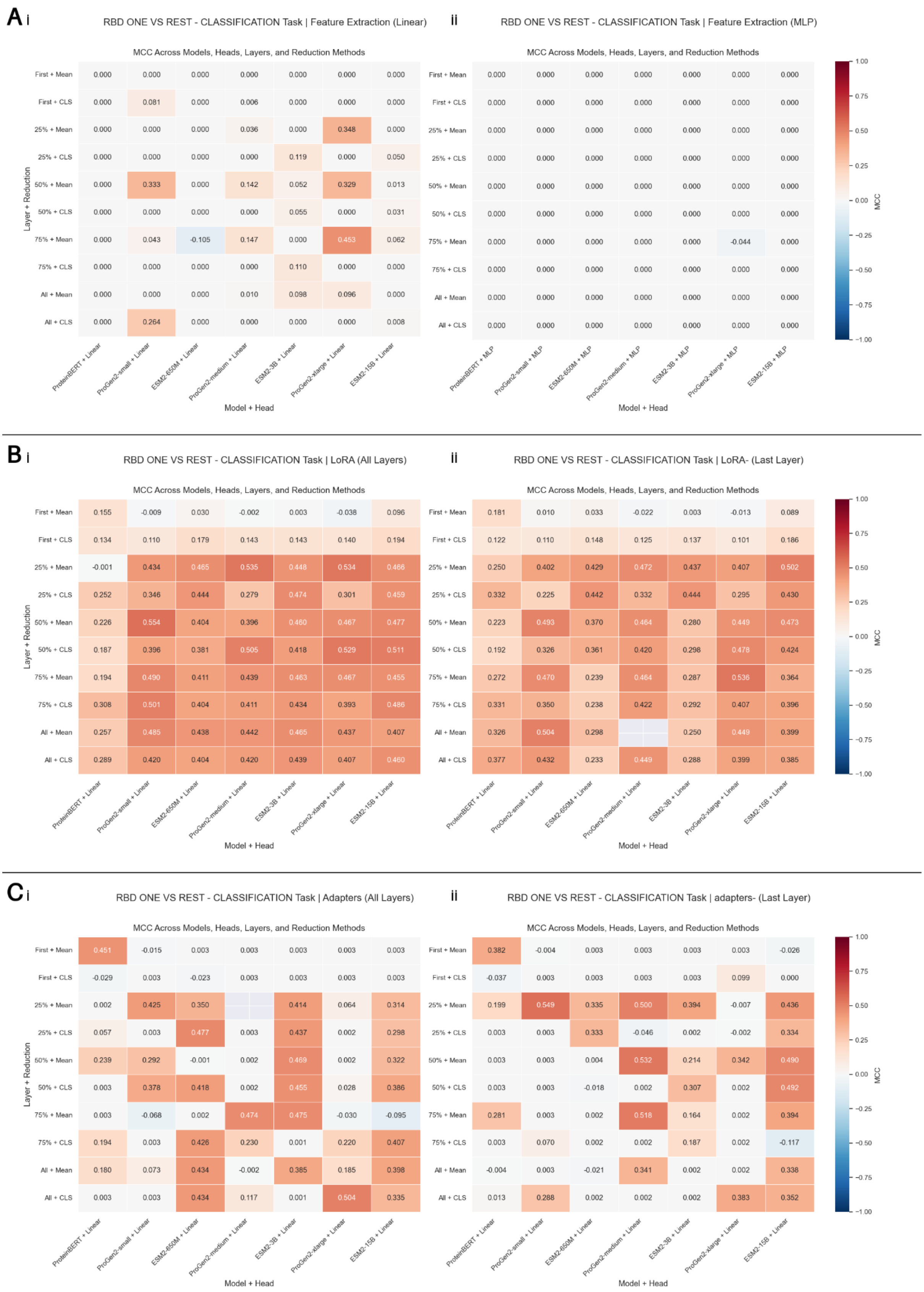


**Figure S6. Detailed results for *RBD-three vs. rest* task.**

MCC is used as a performance metric for each setup, with x-axis showing the PLM and the head used and the y-axis representing the layers used and the pooling method employed; mean stands for mean pooling and CLS for pooling the classification token for BERT-based PLMs (ESM2, ProteinBERT) and the EOS token for GPT-based PLMs (ProGen2). (A) Feature extraction detailed results using (i) a linear downstream head and (ii) a MLP with one hidden layer as a downstream head. (B) LoRA detailed results when (i) applying LoRA to all layers of PLMs and (ii) applying LoRA to the last layer of PLMs. (C) Adapters detailed results when

(i) applying adapters to all layers of PLMs and (ii) applying adapters to the last layer of PLMs. Empty cells represent work in progress, due to the computational burden these setups bear.

TL: Transfer Learning; PLM: Protein Language Model; LoRA: Low Rank Adaptation

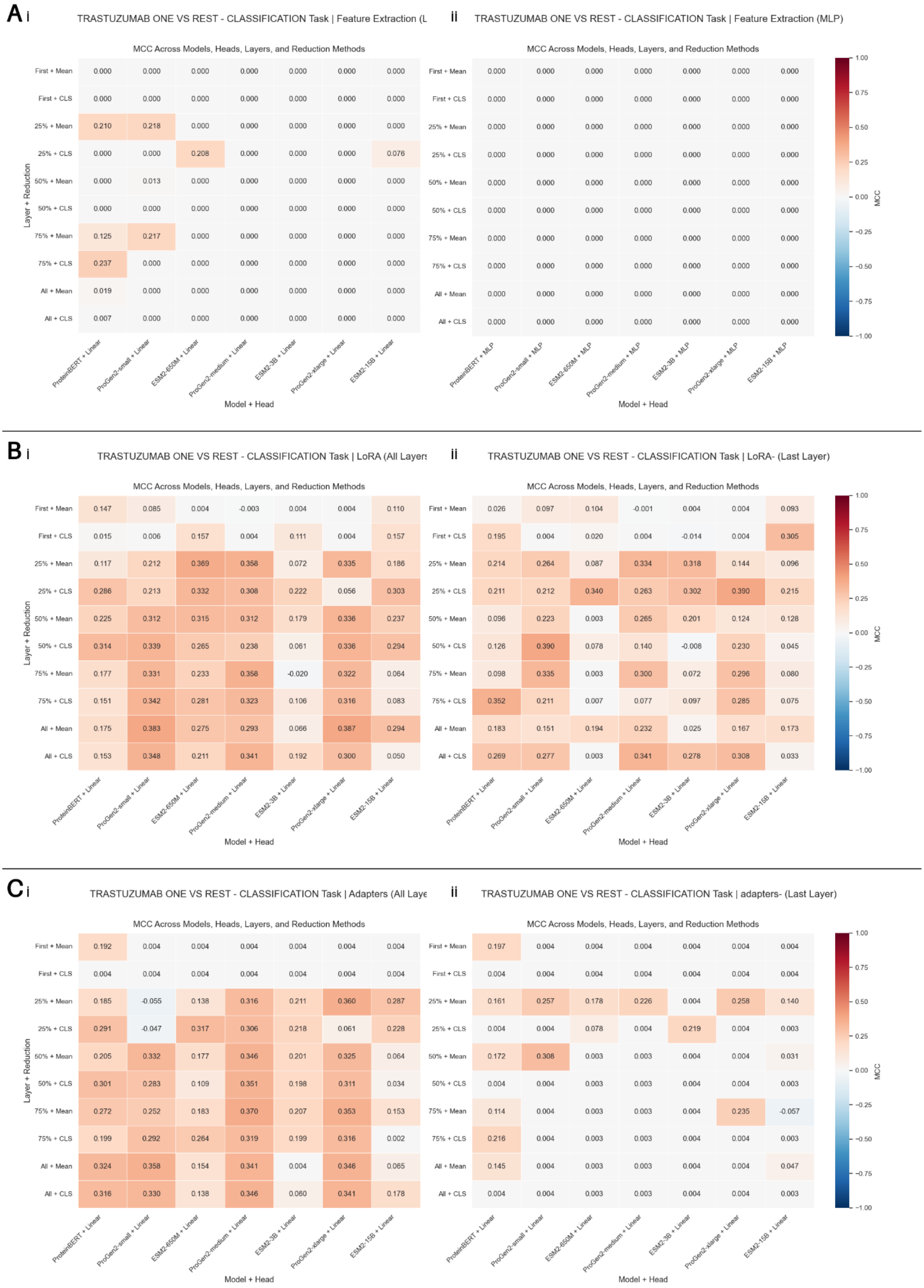


**Figure S7. Detailed results for *Trastuzumab-three vs. rest* task.**

MCC is used as a performance metric for each setup, with x-axis showing the PLM and the head used and the y-axis representing the layers used and the pooling method employed; mean stands for mean pooling and CLS for pooling the classification token for BERT-based PLMs (ESM2, ProteinBERT) and the EOS token for GPT-based PLMs (ProGen2). (A) Feature extraction detailed results using (i) a linear downstream head and (ii) a MLP with one hidden layer as a downstream head. (B) LoRA detailed results when (i) applying LoRA to all layers of PLMs and (ii) applying LoRA to the last layer of PLMs. (C) Adapters detailed results when

(i) applying adapters to all layers of PLMs and (ii) applying adapters to the last layer of PLMs. Empty cells represent work in progress, due to the computational burden these setups bear.

TL: Transfer Learning; PLM: Protein Language Model; LoRA: Low Rank Adaptation

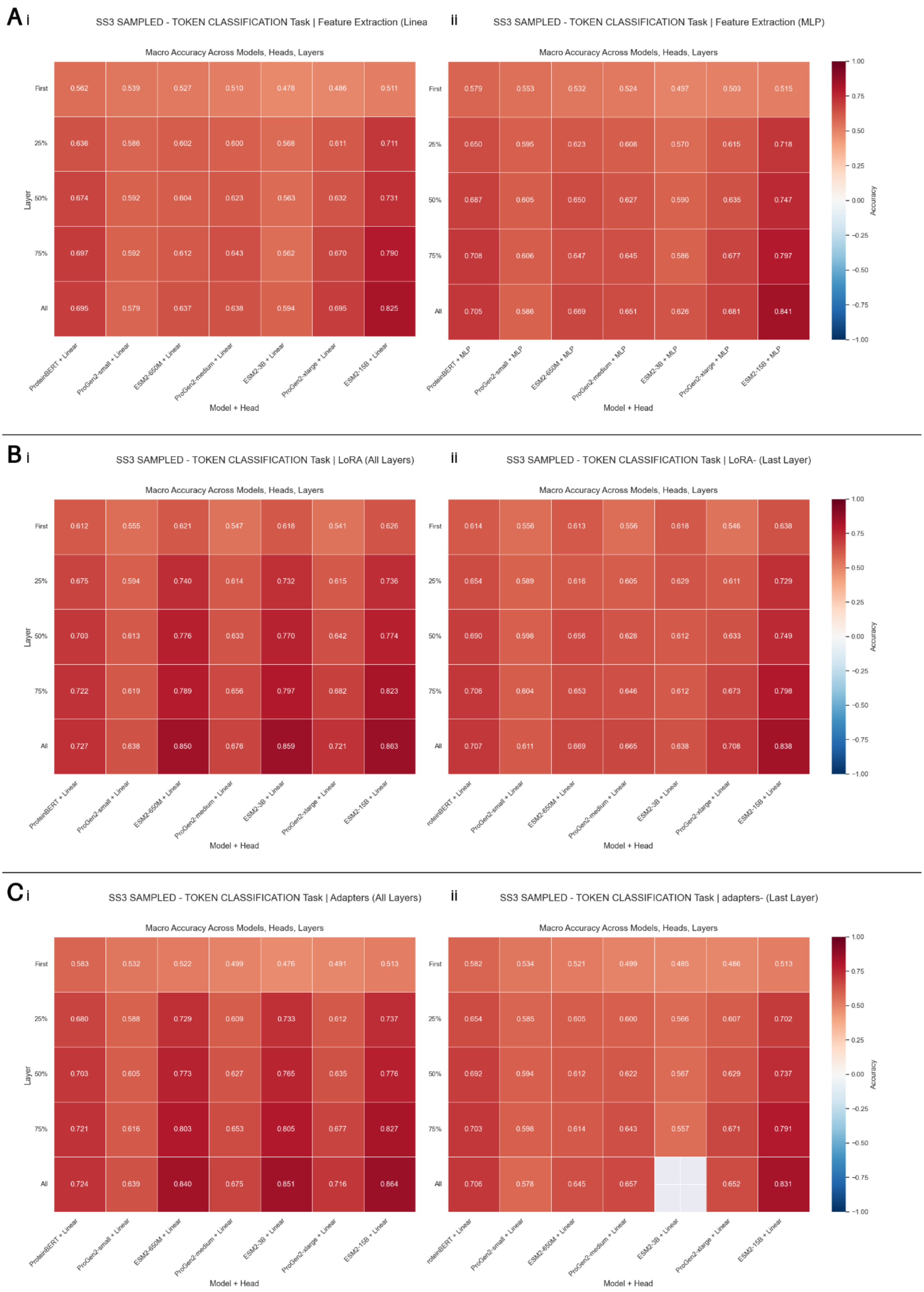


**Figure S8. Detailed results for *SS3-sampled* task.**

Spearman's rank correlation is used as a performance metric for each setup, with x-axis showing the PLM and the head used and the y-axis representing the layers used; no pooling was performed. (A) Feature extraction detailed results using (i) a linear downstream head and (ii) a MLP with one hidden layer as a downstream head. (B) LoRA detailed results when (i) applying LoRA to all layers of PLMs and (ii) applying LoRA to the last layer of PLMs. (C) Adapters detailed results when (i) applying adapters to all layers of PLMs and (ii) applying adapters to the last layer of PLMs. Empty cells represent work in progress, due to the computational burden these setups bear.

TL: Transfer Learning; PLM: Protein Language Model; LoRA: Low Rank Adaptation
